# Novel 4D tensor decomposition-based approach integrating tri-omics profiling data can identify functionally relevant gene clusters

**DOI:** 10.64898/2026.03.19.712900

**Authors:** Turki Turki, Y.-H. Taguchi

## Abstract

Understanding gene expression requires integrating multiple regulatory layers, because transcript abundance does not necessarily correspond to translational activity or protein abundance. Ribosome profiling and proteomics help distinguish increased translation from ribosome stacking or translational buffering, but no de facto standard framework exists for unsupervised integration of transcriptome, translatome, and proteome profiles. Here, we propose a four-dimensional tensor decomposition-based unsupervised feature extraction approach for tri-omics integration. We applied higher-order singular value decomposition to transcriptome, Ribo-seq, and proteome profiles measured under branched-chain amino acid starvation. The resulting singular value vectors captured relationships among the three omics layers, including a component consistent with ribosome stacking, where transcrip-tome and translatome signals increased while proteome signals decreased, and another consistent with translational buffering, where proteome variation was suppressed despite transcriptome and translatome changes. Gene selection identified 1,781 genes associated with ribosome stacking and 227 genes associated with translational buffering. Enrichment analyses linked the former to translation, post-translational protein modification, RNA polymerase II transcription, cell cycle regulation, endoplasmic reticulum protein processing, ubiquitin-mediated proteolysis, and stress-related pathways, and the latter to ribosome, translation elongation and termination, spliceosome, immune- and stress-related pathways, and ribosomopathy-associated diseases. Robustness analyses indicated that the results were not substantially affected by the duplicated proteome replicate or missing-value handling. Under the tested settings, comparison with MOFA+ and mixOmics suggested that our approach more directly extracted components interpretable as ribosome stacking and translational buffering. These results demonstrate that tensor decomposition-based unsupervised feature extraction is useful for identifying functionally relevant gene clusters from tri-omics data.

## 1. Introduction

Despite its apparent ease, measuring gene expression is not straightforward. The purpose of measuring gene expression is to identify genes that are active in a context-dependent manner. Relating the phenotype to the amount of expressed genes can help us to infer the functions of the genes. Thus, it is important to assume that more highly expressed genes are more active in some contexts. Nevertheless, expression levels are not always related to the activity of individual genes. This is because the definition of gene expression remains unclear. It is typical to measure gene expression as the amount of mRNA. However, the amount of mRNA is not always equivalent to the activity of the genes, because mRNA cannot function until it is translated into a protein. One might wonder why we measure the amount of mRNA rather than directly measuring the amount of protein. Defining gene expression as the amount of protein creates another problem, as the amount of protein is not directly regulated [1]. Most of the molecular machinery that can influence gene expression is in mRNA layer, e.g. promoter methylation, protein binding to DNA, or non-coding RNA that can affect the amount of mRNA after its transcription from DNA. Thus, if we define protein expression as gene expression, we cannot directly relate the molecular machinery to the amount of gene expression. Furthermore, the amount of mRNA is not always directly related to the amount of protein present. If translation is halted, the amount of protein will not increase even if mRNA expression increases. In contrast, even if the amount of mRNA remains stable, the amount of protein can increase if the speed of translation increases [2]. In addition, compared to our knowledge of the molecular machinery that regulates the amount of mRNA, we have little knowledge about how the speed of translation is regulated. Translation speed is strongly related to the amount of ribosome. More ribosome usually results in more proteins. Thus, it appears sufficient to measure mRNA, ribosomes, and proteins simultaneously. However, the simultaneous measurement of mRNA, ribosomes, and proteins is not sufficient to determine gene expression in a context-dependent manner. More ribosome does not always increase translation speed, as the increased amount of ribosome could reflect ribosome stacking [3], which is evidence of decreased translation speed. Thus, we need to determine how to distinguish between increased ribosomes that reflect an increased translation speed and those caused by ribosome stacking.

There is a long history of studies related to the interrelationships between mRNA, ribosome profiling (Ribo-seq) [4], and the proteome. Hereafter, we refer to these as the transcriptome, translatome, and proteome. Initially, Schwanhäusser et al. [5] proposed that as little as 40 percent of gene expression is regulated at the transcriptome level, whereas as much as 60 percent is regulated at the proteome level. Nevertheless, based on tri-omics (transcriptome, translatome, and proteome) [6], Jovanovic et al. [7] found that variations in the proteome are usually regulated at the transcriptome level, at least for steady-state and dynamic responses. In contrast, proteome modeling (changes in cell function) is governed directly at the proteome level. Thus, translational buffering (TB) [8] is a central issue in tri-omics studies.

In 2015, Battle et al. [9] and Cenik et al. [10] proposed the concept of TB, referring to the suppression of proteome variation despite transcriptome variation owing to control by the translatome layer. This mechanism is useful to maintain proteome stability during transcriptome variation. Two possible mechanisms underly TB:

- Translation initiation rate model: as mRNA levels increase, translation initiation factors and ribosomes become rate-limiting, reducing the frequency of translation initiation per mRNA molecule. Consequently, even with increased mRNA, total protein synthesis reaches saturation.
- Differential accessibility model: excess mRNA is sequestered into pools not utilized for translation (P-bodies, stress granules, or specific ribonucleoprotein complexes). This regulates the proportion of translatable mRNA (Fraction), altering the apparent translation efficiency.

Several studies have investigated this topic. Blevins et al [11] found the followings

- The proteome is more strongly correlated with the translatome than the transcriptome.
- TB occurs broadly.
- Transcriptome suppression plays a critical role in some gene groups (e.g., cell cycle-regulated genes).

Tri-omics have also been analyzed at the isoform level [12] but better proteome accuracy is required for a better understanding. Lu et al. [13] found that the high correlation between the translatome and proteome could be destroyed in some specific cases. Cuevas et al [14] considered a non-canonical proteome in tri-omics analysis. Zu et al. [15] considered plant tri-omics and confirmed that the proteome is more strongly correlated with the translatome than the transcriptome. Worpenberg et al. [16] investigated BCAA starvation in tri-omics and determined the importance of the dependence of dwell time on the condition. Elpida et al. [17] also investigated TB using tri-omics.

Despite these numerous studies using tri-omics (transcriptome, translatome, and proteome) a *de facto* standard method to integrate tri-omics does not appear to exist. We propose the use of tensor decomposition (TD)-based unsupervised feature extraction (FE) [18]. TD-based unsupervised FE has been successfully analyzed in numerous multiomics integration studies; therefore, it would not be surprising if it could be used to successfully integrate tri-omics. To determine whether TD-based unsupervised FE can successfully integrate tri-omics, we applied it to the datasets of Worpenberg et al. [16], which contain the largest number of samples among datasets associated with tri-omics measurements. As a result, we showed that TD-based unsupervised FE successfully identified functional gene sets associated with high and low translational efficiency and identified their functional mechanisms. Thus, we can conclude that TD-based unsupervised FE can satisfactorily integrate tri-omics.

These studies [19–23] demonstrate the growing importance of deep representation learning, graph-based modeling, and foundation models. However, our aim is different: interpretable unsupervised integration of bulk transcriptome, Ribo-seq, and proteome data with explicit omics-mode loadings.

## 2. Materials and Methods

### 2.1. Datasets

The datasets analyzed in this study were generated by Worpenberg et al. [16], who investigated codon-specific ribosome stalling during branched-chain amino acid starvation in NIH3T3 mouse fibroblast cells. Cells were analyzed under six conditions: complete medium control, leucine starvation, isoleucine starvation, valine starvation, double starvation of leucine and isoleucine, and triple starvation of leucine, isoleucine, and valine. RNA-seq and Ribo-seq data were obtained from matched experimental conditions, and quantitative proteomics data were obtained from the same starvation design. These datasets were used to construct the tri-omics tensor described in Section 2.2

#### 2.1.1. Gene expression profiles

Gene expression profiles were retrieved from GEO with GEO ID GSE291652. The supplementary file, “GSE291652_RNA_seq_raw_count.tsv.gz”, was downloaded from GEO and loaded into R using the `read.csv` command.

#### 2.1.2. Ribo-seq profiles

Ribo-seq expression profiles were retrieved from GEO using the GEO ID GSE291653. The supplementary file, “GSE291653_Ribo_seq_raw_count.tsv.gz”, was downloaded from GEO and loaded into R using the `read.csv` command.

#### 2.1.3. Proteome

The Proteome, archived in PRIDE with ID PXD067949 [24], was in sheet 4 of “13059_2025_3800_MOESM4_ESM.xlsx” obtained from supplementary file [16]; it was loaded into R using the `read_excel` command.

### 2.2. Tensor formation from three omics profiles

Three omics profiles (gene expression, the Ribo-seq profile, and the proteome) were formatted as tensor *x*_*ijmk*_ ∈ ℝ^*N*×6×3×3^, which represents the *k*th omics profile (*k* = 1: gene expression, *k* = 2: Ribo-seq, and *k* = 3: proteome) was attributed to *i*th gene measured under the *j*th experimental condition (*j* = 1:control, *j* = 2:Leu starvation, *j* = 3:Ile starvation, *j* = 4:Val starvation, *j* = 5: double (Leu and Ile) starvation, and *j* = 6: triple (all) starvation) of the *m*th replicate. *N* = 18, 175 is the number of genes in the gene expression and Ribo-seq profiles. During this process, because there were only two replicates of proteome for *j* = 5 (double starvation), *m* = 1 was replicated (doubled). The number of detected proteins (7, 004) was less than *N*(=18,175). *x*_*ijmk*_ for missing proteins was filled with zero. Proteins whose corresponding genes were not expressed in the gene or Ribo-seq datasets were discarded. Gene symbols corresponding to proteins were converted to Ensembl gene IDs, employed by the gene expression and Ribo-seq datasets, using bioMart [25], and merged with gene expression and Ribo-seq profiles. *x*_*ijmk*_ is logarithmically transformed as

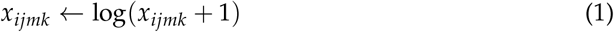

1 is added to prevent divergence when *x*_*ijmk*_ = 0.

### 2.3. Tensor decomposition-based unsupervised feature extraction

Here, we briefly introduce TD-based unsupervised FE. Further details can be found in [18]. After *x*_*ijmk*_ is standardized,

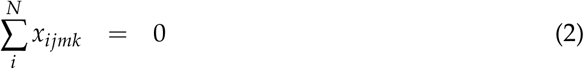

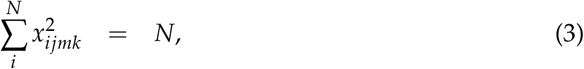

higher-order singular value decomposition (HOSVD) [18] is applied to *x*_*ijmk*_ to obtain

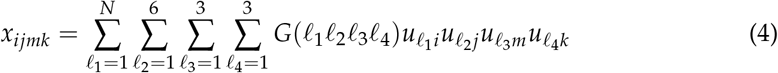

where *G* ∈ ℝ^*N*×6×3×3^ is a core tensor that represents the contribution of 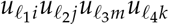 toward *x*_*ijmk*_, and 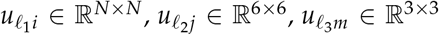, and 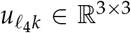 are singular value matrices and orthogonal matrices.

In TD-based unsupervised FE, we select genes using 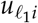 Prior to identifying 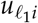 used for gene selection, we must specify that where 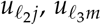 and 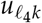 are the parameters of interest. First, 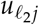 should be distinct between the control (*j* = 1) and others (2 ≤ *j* ≤ 6). 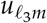 should be constant because the *m*s are replicates. What is expected for 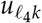 is highly context dependent. We must specify the 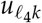 of interest only after investigating the computed 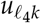.

Once we identify which 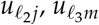 and 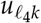 are of interest, we investigate the absolute value of *G* using the fixed values of *ℓ*_2_, *ℓ*_3_, and *ℓ*_4_. 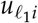 whose *ℓ*_1_ was associated with the largest absolute G, was used to select the genes. The *P*-value, *P*_*i*_, is attributed to the *i*th gene, assuming that 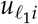 obeys the Gaussian (empirical null hypothesis)

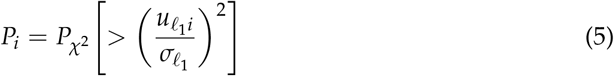

where 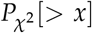 is the cumulative *χ*^2^ distribution in which the argument is larger than *x* and 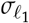 is the standard deviation.

Here, 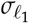 is optimized as follows:

1. Set the initial value of 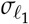.
2. Assign *P*_*i*_ to *i*s with eq. (5).
3. Collect *P*_*i*_ using BH criterion [18].
4. Exclude *i*s with adjusted *P*-values less than the threshold value, 0.01.
5. Compute the histogram *h*_*s*_(1 − *P*_*i*_), 1 ≤ *s* ≤ *S*.
6. Compute the standard deviation, 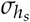, of *h*_*s*_.
7. Modify 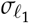 such that 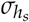 decreases.
8. Go back to step 2 if 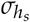 is not minimized, otherwise terminate the process.

Subsequently, *P*_*i*_ is reassigned to *i* using eq. (5) and *i*s with adjusted *P*-values less than 0.01 are selected.

### 2.4. Enrichment analysis

Enrichment analysis was performed with uploading the set of selected either 1,781 or 227 gene symbols to either DAVID [26] or Enrichr [27] (Table 1). Results from DAVID and

**Table 1.**
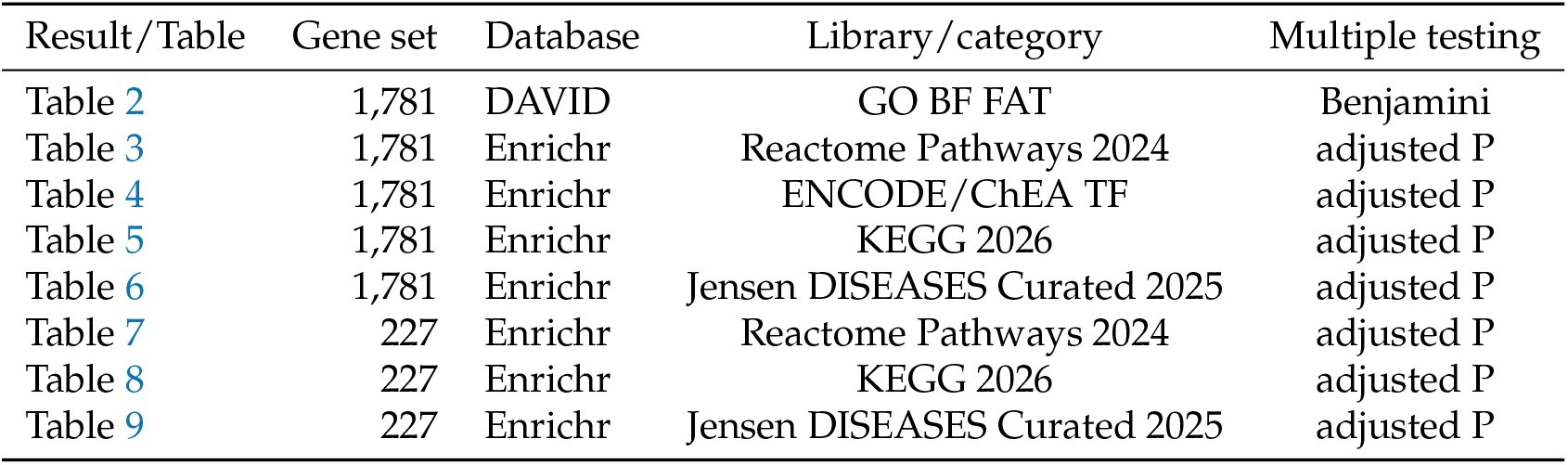
Summary of enrichment analysis results.

| Result/Table | Gene set | Database | Library/category | Multiple testing |
| --- | --- | --- | --- | --- |
| Table 2 | 1,781 | DAVID | GO BF FAT | Benjamini |
| Table 3 | 1,781 | Enrichr | Reactome Pathways 2024 | adjusted P |
| Table 4 | 1,781 | Enrichr | ENCODE/ChEA TF | adjusted P |
| Table 5 | 1,781 | Enrichr | KEGG 2026 | adjusted P |
| Table 6 | 1,781 | Enrichr | Jensen DISEASES Curated 2025 | adjusted P |
| Table 7 | 227 | Enrichr | Reactome Pathways 2024 | adjusted P |
| Table 8 | 227 | Enrichr | KEGG 2026 | adjusted P |
| Table 9 | 227 | Enrichr | Jensen DISEASES Curated 2025 | adjusted P |

Enrichr were not merged; each table reports the output from the specified database/library.

### 2.5. Using generative AI to understand the biological meaning of the gene sets

Because the number of genes selected was large, we needed to understand the biological significance of the selected genes. Enrichment analysis is the best method for achieving this goal. Nevertheless, there were too many enriched terms among too many categories, making it difficult to understand the biological significance of the selected genes based only on enrichment analysis. Therefore, we used generative AI.

#### 2.5.1. Direct analysis using generative AI

All the sets of genes were uploaded to Gemini Pro 3.0 Deep Research with the script “Please consider what functional units exist within this subset of genes” in Japanese. The summaries were individually evaluated using a manual literature search. Only those evaluated in the manual literature search were included in this study. An infographic of the summary was also generated with the script “Please create an infographic illustrating the relationships between these gene subunits” in Japanese.

#### 2.5.2. Analysis using generative AI after clustering enrichment analysis

All the sets of genes were loaded into DAVID, and annotations were clustered using the function implemented in DAVID [26]. The list of clustered annotations was uploaded to Gemini Pro 3.0 Deep Research with the script “Please consider what functional units exist within this subset of genes” in Japanese. The summaries were individually evaluated using a manual literature search. Only those evaluated in the manual literature search were included in this study. An infographic of the summary was also generated with the script “Please create an infographic illustrating the relationships between these clusters” in Japanese.

### 2.6. Comparison with state-of-the-art (SOTA)

For a fair comparison, MOFA+ and mixOmics were applied to the same preprocessed tri-omics data used immediately before applying TD-based unsupervised FE. The pre-processing procedure, including gene matching, log(x + 1) transformation, missing-value handling, and treatment of the duplicated proteome replicate, was the same as described in Section 2.2. Because MOFA+ and mixOmics require matrix-form input rather than a four-dimensional tensor, the preprocessed tensor was converted into omics-specific matrices with matched condition–replicate samples.

#### 2.6.1. MOFA+

To evaluate whether MOFA+ could recover proteome-associated latent factors, we ran MOFA+ with settings intended to reduce premature factor dropping and to allow weak proteome-associated signals to be retained.

~~~
# Increase the number of factors to detect minor contribution model_opts$num_factors <-10
# Learning Option: Relax ARD (Factor Selection) train_opts <-get_default_training_options(MOFAobject)
# Increase the number of training iterations
# so that even very small changes are not ignored
train_opts$maxiter <-2000
train_opts$convergence_mode <- “slow”
train_opts$drop_factor_threshold <-0
# [Important] Execute this function separately before running run_mofa MOFAobject <-prepare_mofa(MOFAobject,
     model_options = model_opts,
     training_options = train_opts)
~~~

The resulting variance explained by MOFA+ factors is reported in Fig. 9 and Table S1. Full code is available on GitHub.

#### 2.6.2. mixOmics

mixOmics is performed with the default setting (attributing the same weight to all pairs in design matrix). Full code is available on GitHub.

### 2.7. Ribosome stacking score and translational buffering score

Because the aim of this score analysis was to evaluate whether the selected genes occupy higher-scoring regions of the full four-dimensional tensor, we did not average the scores across conditions or replicates. Averaging across experimental conditions would collapse biologically distinct starvation responses. Therefore, the Wilcoxon rank-sum test was used as a tensor-entry-level distributional comparison over gene–condition–replicate scores, rather than as a gene-level test based on one averaged score per gene.

#### 2.7.1. Ribosome stacking score

To compute ribosome stacking score corresponding to eq. (8), we compute

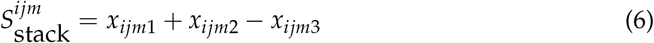

and apply Wilcoxon rank-sum test between {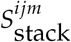 | 1,781 genes} and {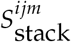 | remaining genes} with the alternative hypothesis {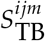 | 1,781 genes} is greater than {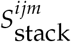 | remaining genes}. Wilcoxon rank-sum test was performed with the function, `wilcox.test`, in R. Because of eqs. (2) and (3), *x*_*ijmk*_ is essentially *z* score.

#### 2.7.2. Translational buffering score

To compute translational buffering score corresponding to eq. (9), we compute

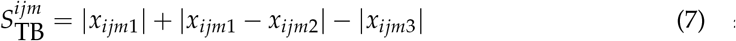

and apply Wilcoxon rank-sum test between {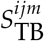 | 227 genes} and {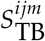 | remaining genes} with the alternative hypothesis {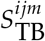 | 227 genes} is greater than {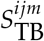 | remaining genes}. Wilcoxon rank-sum test was performed with the function, `wilcox.test`, in R. Because of eqs. (2) and (3), *x*_*ijmk*_ is essentially *z* score.

### 2.8. Computational complexity / scalability

TD-based unsupervised FE required limited computational resources in the present dataset. In the current setup, the tensor contained 18,175 × 6 × 3 × 3 = 981,450 entries. Runtime was approximately 10 seconds on 12th Intel(R) Core(TM) i7-1270P, with peak memory below 32 GB. As for the scalability, HOSVD/SVD can be applied to larger problems by using truncated SVD or irlba.

### 2.9. Limitations

This study does not provide direct experimental validation of ribosome stacking or translational buffering. The proposed interpretation is based on latent component patterns, enrichment analysis, and orthogonal computational evidence. Experimental perturbation of selected genes and direct measurement of ribosome collision or protein synthesis will be required in future studies.

## 3. Results

### 3.1. Investigation of singular value vectors

First, we derive the singular value vectors 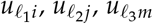, and 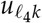 by applying TD to *x*_*ijmk*_. Among them, we consider 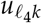 because it is important to check the relationships among the tri-omics, represented by 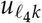 where *k* represents the dependence on individual tri-omics (Fig. 1). Among 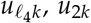 is interesting because *u*_23_, which represents the proteome, decreases, whereas *u*_21_, which represents the transcriptome, and *u*_22_, which represents the translatome, increase. This suggests that ribosome stacking occurs because the proteome decreases despite the increased transcriptome and translatome. In addition, *u*_3*k*_ is interesting because *u*_33_, which represents the proteome does not change, whereas *u*_31_, which represents the transcriptome, increases and *u*_32_, which represents the translatome, decreases. This suggests that *u*_3*k*_ represents TB.

**Figure 1.**
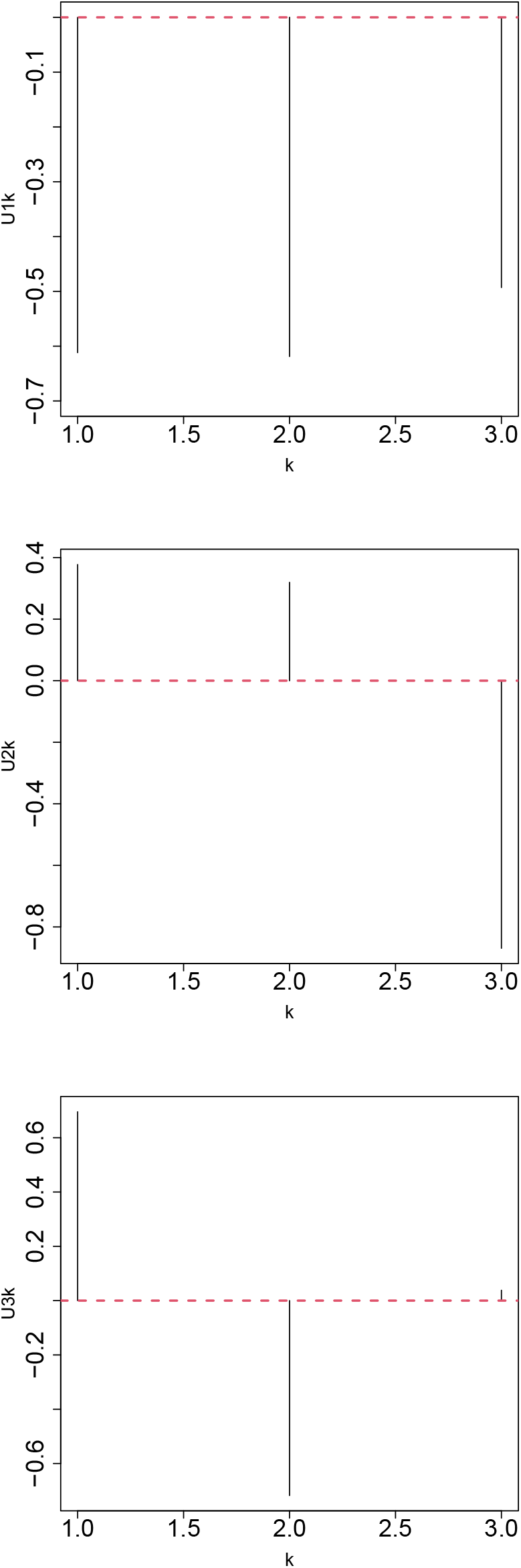
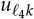 top: *ℓ*_4_ = 1, middle: *ℓ*_4_ = 2. bottom: *ℓ*_4_ = 3. *k* = 1: transcriptome, *k* = 2: translatome, *k* = 3: proteome

One might wonder whether regarding *u*_2*k*_ and *u*_3*k*_ as ribosome stacking and TB, respectively, is reasonable or not, since it might be overinterpretation without the direct evidences. Nevertheless, it is rare to get vectors consistent with ribosome stacking and TB, since even MOFA+ [28] as well as mixOmics [29], which are two leading SOTA for multiomics integration, did not recover comparable tri-omics components under the tested settings (see below).

In the following section, we consider *u*_23_ because ribosome stacking is likely to contribute to more biological processes than TB, which simply maintains the stability of the proteome.

However, 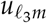 should be independent of *m* and *ℓ*_3_ = 1 satisfies this requirement (not shown here).

### 3.2. *Gene selection: ℓ*_1_ = 6

Subsequently, to determine whether 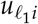 or 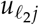 should be considered for ribosome stacking, we investigated *G*(*ℓ*_1_, *ℓ*_2_, 1, 2) (Fig. 2). It is clear that (*ℓ*_1_, *ℓ*_2_) = (2, 1) is the largest, and (*ℓ*_1_, *ℓ*_2_) = (6, 2) is the second largest. To determine which is valid, we must compare *u*_1*j*_ with *u*_2,*j*_ (Fig. 3). Because *u*_1*j*_ does not exhibit any *j*-dependence, *u*_2*j*_ should be considered. Consequently, *u*_6*i*_, which shares a large *G* with *u*_2*j*_ and *u*_2*k*_, should be used for gene selection. The left panel of Fig. 4 represents the dependence of the standard deviation of the histogram of 1 − *P*_*i*_ upon 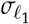. It is obvious that the standard deviation of the histogram of 1 − *P*_*i*_ reaches a minimum at some value of 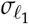. *P*_*i*_ is recomputed using the estimated value of 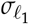. The right panel of Fig. 4 shows the histogram. The large peak on the right side corresponds to the selected genes. *P*-values were corrected using the BH criterion [18]. The 1,781 genes with adjusted *P*-values less than 0.01 are selected and used for further analysis.

**Figure 2.**
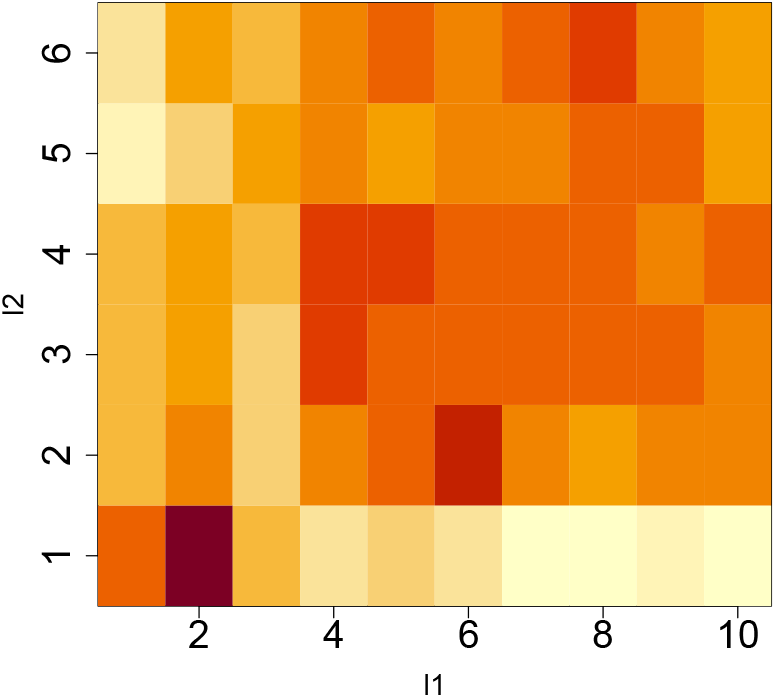
Logarithmic absolute value of *G*(*ℓ*_1_ *ℓ*_2_, 1, 2), log |*G*(*ℓ*_1_, *ℓ*_2_, 1, 2)|. Horizontal:*ℓ*_1_ and vertical: *ℓ*_2_. Darker colors indicate larger values.

**Figure 3.**
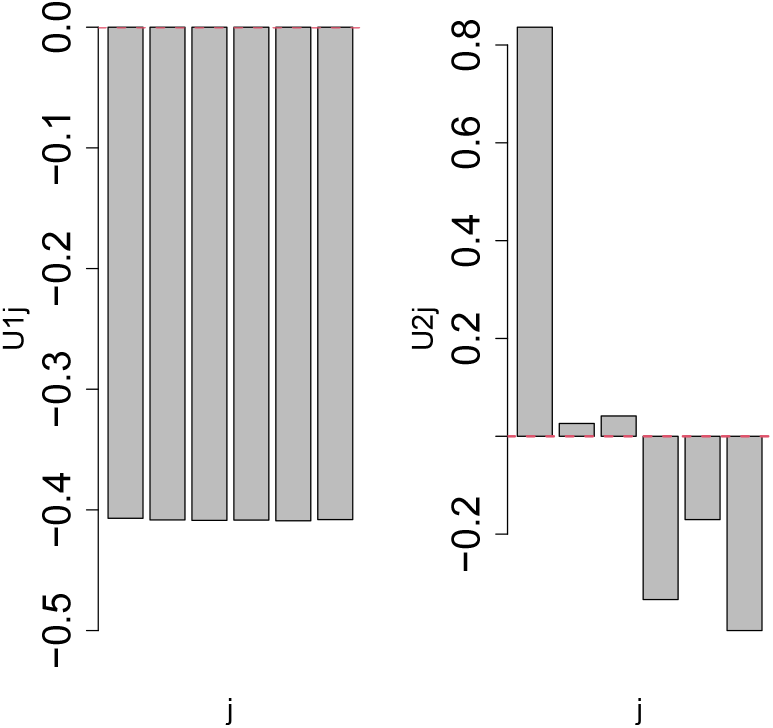
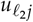 left: *ℓ*_2_ = 1, right: *ℓ*_2_ = 2. *j* = 1:control, *j* = 2:Leu starvation, *j* = 3:Ile starvation, *j* = 4:Val starvation, *j* = 5: Double (Leu and Ile) starvation, and *j* = 6: Triple (all) starvation.

**Figure 4.**
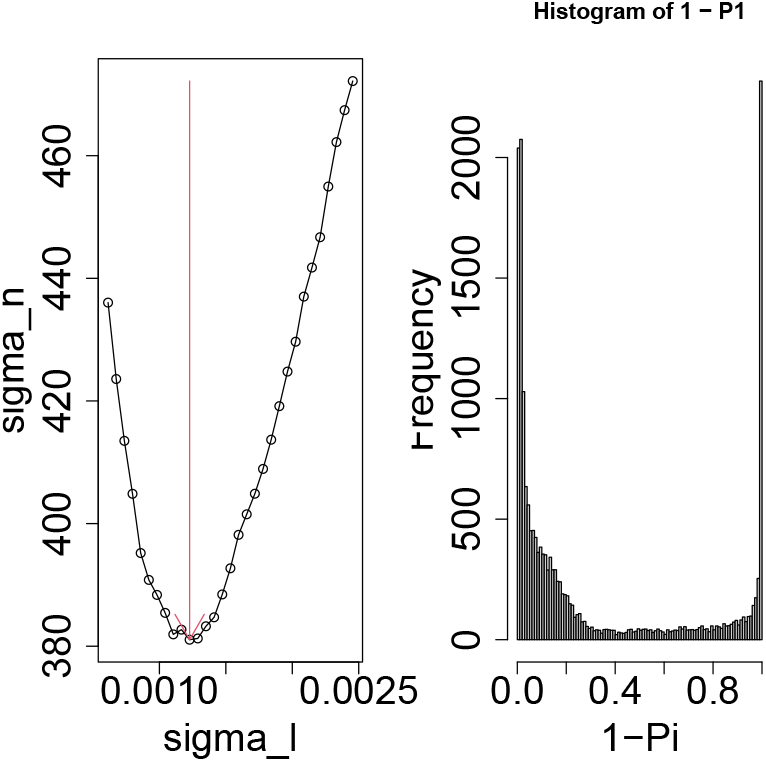
Left: Dependence of the standard deviation (vertical axis) of the histogram of 1 − *P*_*i*_ on 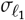 (horizontal axis). Vertical red arrow indicates the estimated 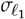. Right: Histogram of 1 − *P*_*i*_ computed using the estimated 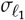.

### 3.3. Validation of ribosome stacking score

To see whether the selected 1,781 genes are associated with ribosome stacking, we define ribosome stacking score as

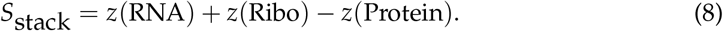

See more details for Methods. When the analysis was restricted to genes with quantified proteome measurements, no significant difference was detected between the 1,781 selected genes and the remaining genes. This restriction excludes genes outside the proteome coverage, many of which are likely to correspond to low-abundance proteins. This does not mean that missing protein measurements were interpreted as true zero protein abundance. Rather, this score uses the same zero-filled proteome layer as that used for tensor construction and should be regarded as a missingness-aware tensor-based score. We therefore computed *S*_stack_ for all genes and investigated whether the selected 1,781 genes showed larger Sstack values than the remaining genes using the Wilcoxon rank-sum test. This tensor-entry-level comparison showed a strong upward shift of *S*_stack_ for the selected 1,781 genes (Wilcoxon rank-sum test, *P* < 2.2 × 10^−16^). Thus we can conclude that TD-based unsupervised FE successfully selected genes associated with ribosome stacking.

### 3.4. Understanding the biological meaning of the selected genes

To see whether the selected set of 1,781 genes are biologically meaningful, we have uploaded the selected set of 1,781 genes to various enrichment analysis servers. Primarily, since it is expected that the set of genes is enriched with regulation of translation, we checked it by uploading them to “GO BF FAT” category in DAVID. Table 2 lists the terms including “translation” and associated with adjusted *P*-values less than 0.05. Since two terms related to post-translational protein modification is enriched, it is quite reasonable.

**Table 2.** GO biological process enrichment results (GO BF FAT in DAVID) including “translation”.

| Term | Count | List Total | Pop Hits | Pop Total | P-Value | Benjamini | Fold Enrichment |
| --- | --- | --- | --- | --- | --- | --- | --- |
| post-translational protein modification | 138 | 1628 | 921 | 19441 | $1.56 \times 10^{-11}$ | $2.98 \times 10^{-9}$ | 1.79 |
| translation | 73 | 1628 | 462 | 19441 | $1.93 \times 10^{-7}$ | $1.51 \times 10^{-5}$ | 1.89 |
| mitochondrial translation | 23 | 1628 | 110 | 19441 | $9.95 \times 10^{-5}$ | $3.94 \times 10^{-3}$ | 2.50 |
| regulation of post-translational protein modification | 40 | 1628 | 256 | 19441 | $1.88 \times 10^{-4}$ | $6.60 \times 10^{-3}$ | 1.87 |
| negative regulation of post-translational protein modification | 19 | 1628 | 98 | 19441 | $1.18 \times 10^{-3}$ | $3.00 \times 10^{-2}$ | 2.32 |
| cytoplasmic translation | 28 | 1628 | 175 | 19441 | $1.42 \times 10^{-3}$ | $3.44 \times 10^{-2}$ | 1.91 |

We have also uploaded the set of the selected 1,781 genes to “Reactome Pathways 2024” category of Enrichr (Table 3). We can notice that the same number of translation as well as transcription related terms are included. This can be interpreted as follows. When ribosome stacking occurs, transcription is known to be affected to terminate over-production of mRNA [30]. Especially, the fact that two terms “RNA Polymerase II Transcription” and “RNA Polymerase II Transcription Termination” are enriched is really suggestive; this might be the consequence of the termination of transcription in response to ribosome stacking.

**Table 3.**
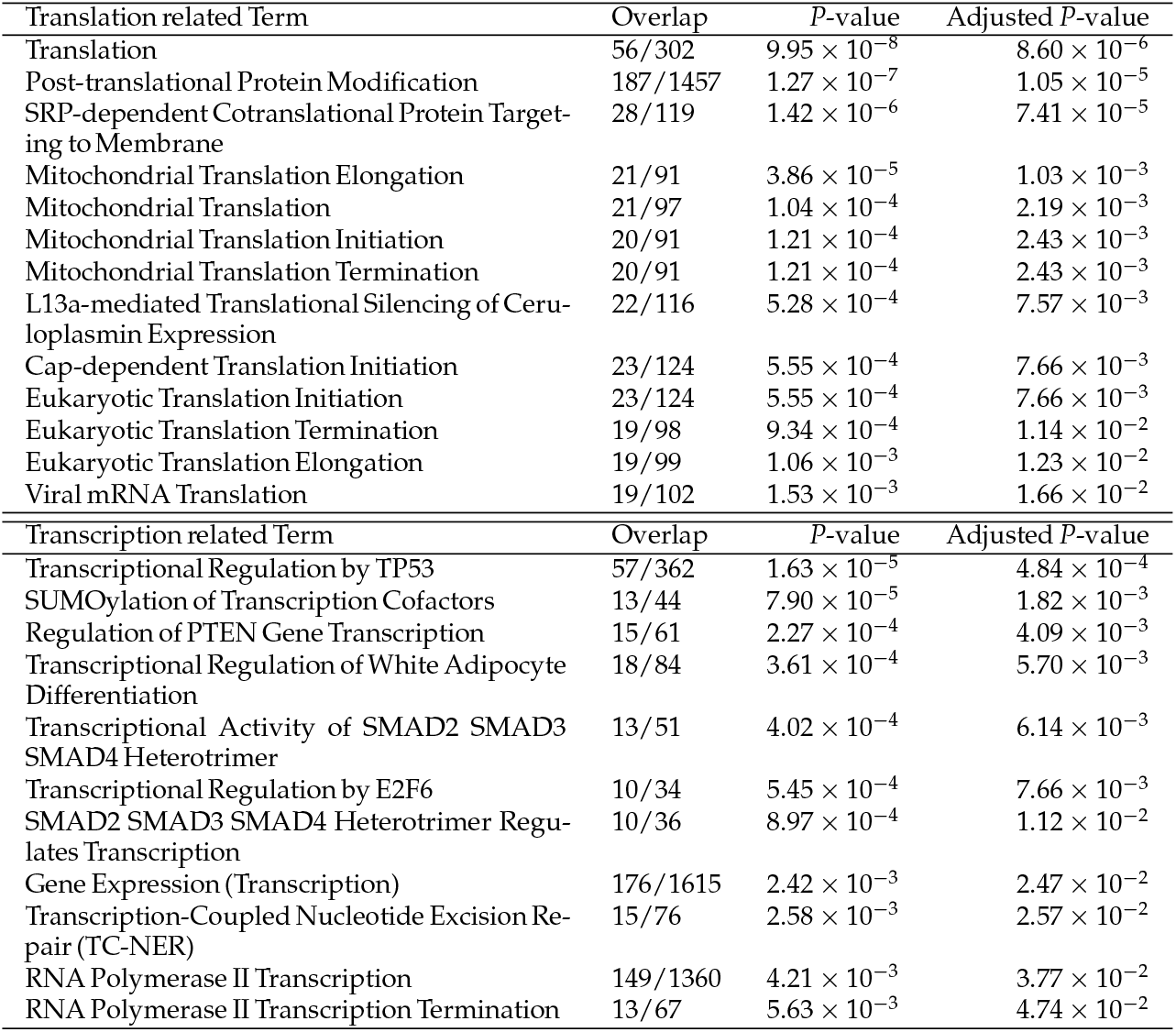
Terms related to “Translation” and “Transcription”, respectively, and associated with adjusted *P*-values less than 0.05 in ‘Reactome Pathways 2024” category of Enrichr.

There is additional evidence that the selected set of 1,781 genes is consistent with the occurrence of ribosome stacking. Table 4 lists the top ranked 10 transcription factor (TF) enrichment in “ENCODE and ChEA Consensus TFs from ChIP-X” category of Enrichr. MYC, MAX, and E2F4, accelerate cell proliferation and protein synthesis [31] and they are downregulated due to energy saving in response to ribosome stacking. ATF2 is known to respond to the ribotoxic stress response (RSR) [32]. TAF1, SIN3A, and YY1, are fundamental factors that regulate transcriptional activity across the entire genome [33–35]; the detection of these TFs suggests that they started to re-organize genomic structure to control transcription in response to ribosome stacking.

**Table 4.** Top ranked 10 transcription factor enrichment in “ENCODE and ChEA Consensus TFs from ChIP-X” category of Enrichr.

| Term | Overlap | *P*-value | Adjusted *P*-value |
| --- | --- | --- | --- |
| TAF1 ENCODE | 491/3346 | $1.04 \times 10^{-33}$ | $1.08 \times 10^{-31}$ |
| NFYB ENCODE | 528/3715 | $7.22 \times 10^{-33}$ | $3.41 \times 10^{-31}$ |
| E2F4 ENCODE | 167/710 | $9.84 \times 10^{-33}$ | $3.41 \times 10^{-31}$ |
| SIN3A ENCODE | 220/1131 | $4.69 \times 10^{-30}$ | $1.22 \times 10^{-28}$ |
| MAX ENCODE | 334/2073 | $3.07 \times 10^{-29}$ | $6.39 \times 10^{-28}$ |
| BRCA1 ENCODE | 454/3218 | $8.45 \times 10^{-27}$ | $1.47 \times 10^{-25}$ |
| ATF2 ENCODE | 405/2852 | $3.96 \times 10^{-24}$ | $5.89 \times 10^{-23}$ |
| E2F6 ENCODE | 443/3245 | $5.21 \times 10^{-23}$ | $6.78 \times 10^{-22}$ |
| MYC ENCODE | 245/1515 | $2.10 \times 10^{-21}$ | $2.43 \times 10^{-20}$ |
| YY1 ENCODE | 384/2753 | $2.58 \times 10^{-21}$ | $2.69 \times 10^{-20}$ |
| E2F1 CHEA | 159/859 | $1.47 \times 10^{-19}$ | $1.39 \times 10^{-18}$ |

Table 5 lists the enriched pathways in “KEGG 2026” category in Enrichr. “CELL CYCLE” and “DNA REPLICATION” are terminated because of ribosome stacking through mTORC1 pathway suppression [36]. The detection of “PROTEIN PROCESSING IN ENDOPLASMIC RETICULUM” is the evidence that ribosomal congestion impairs normal protein folding; this suggests that, particularly in the endoplasmic reticulum (ER)—where membrane proteins and other molecules are synthesized—improperly folded proteins may be accumulating, leading to “endoplasmic reticulum stress.” [37] Nutritional starvation (BCAA deficiency) requires a large-scale reprogramming of gene expression; the “POLYCOMB REPRESSIVE COMPLEX” plays a role in “turning off” specific groups of genes, and this process is thought to be an attempt to silence genes involved in growth in order to adapt to a state of starvation [38]. The “HIPPO SIGNALING PATHWAY” acts as a sensor that regulates cell size and proliferation [39]; it detects physical and chemical stresses, such as ribosomal congestion and amino acid shortages, and functions as a signal that switches the cell into a mode prioritizing survival. The detection of “DNA REPLICATION” may reflect the activity of a backup mechanism that attempts to repair DNA instability caused by factors such as delays in DNA replication. In conclusion, Table 5 can be understood as follows:

**Table 5.** Top ranked 10 pathway enrichment results in “KEGG 2026” category in Enrichr.

| Term | Overlap | *P*-value | Adjusted *P*-value |
| --- | --- | --- | --- |
| RIBOSOME | 37/160 | $4.99 \times 10^{-8}$ | $1.55 \times 10^{-5}$ |
| CELL CYCLE | 36/157 | $9.47 \times 10^{-8}$ | $1.55 \times 10^{-5}$ |
| PROTEIN PROCESSING IN ENDOPLASMIC RETICULUM | 33/171 | $1.72 \times 10^{-5}$ | $1.48 \times 10^{-3}$ |
| UBIQUITIN MEDIATED PROTEOLYSIS | 29/142 | $1.81 \times 10^{-5}$ | $1.48 \times 10^{-3}$ |
| POLYCOMB REPRESSIVE COMPLEX | 19/82 | $8.38 \times 10^{-5}$ | $5.29 \times 10^{-3}$ |
| HIPPO SIGNALING PATHWAY | 29/155 | $9.68 \times 10^{-5}$ | $5.29 \times 10^{-3}$ |
| DNA REPLICATION | 11/35 | $1.51 \times 10^{-4}$ | $5.98 \times 10^{-3}$ |
| HOMOLOGOUS RECOMBINATION | 12/41 | $1.64 \times 10^{-4}$ | $5.98 \times 10^{-3}$ |
| AGE-RAGE SIGNALING PATHWAY IN DIABETIC COMPLICATIONS | 21/100 | $1.64 \times 10^{-4}$ | $5.98 \times 10^{-3}$ |
| GLYCEROPHOSPHOLIPID METABOLISM | 21/102 | $2.19 \times 10^{-4}$ | $7.20 \times 10^{-3}$ |

- Cause: Insufficient BCAAs lead to ribosomal congestion (“RIBOSOME”).
- Result: Protein synthesis is disrupted, resulting in the production of waste (incomplete proteins) (“PROTEIN PROCESSING IN ENDOPLASMIC RETICULUM” and “UBIQUITIN MEDIATED PROTEOLYSIS”).
- Response: Cell proliferation is halted (“CELL CYCLE” and “DNA REPLICATION”), the cell switches to energy-saving mode at the genetic level (“POLYCOMB REPRESSIVE COMPLEX”), and survival signals are emitted (“HIPPO SIGNALING PATHWAY”).

Table 6 lists the top 10 diseases in “Jensen DISEASES Curated 2025” of Enrichr. Metabolic diseases are known to be related to BCAA starvation (“DISEASE”, “INHERITED METABOLIC DISORDER”, and “INHERITED METABOLIC DISORDER”) [40]. Ribosomal stalling and collisions tend to worsen in the presence of specific genetic mutations, and these findings indicate that the “group of genes sensitive to BCAA deprivation” overlaps with the causative genes of many known hereditary diseases (“GENETIC DISEASE”) [41]. Defects in protein synthesis caused by ribosome stacking are likely ranked highly because they can easily damage the structural integrity of specific tissues (“DISEASE OF ANATOMICAL ENTITY”), such as the musculoskeletal system and blood vessels [42]. BCAAs are a source of nutrition for cancer cells (“CANCER”) [43]. Rapidly proliferating cells (such as cancer cells) have a high rate of translation and are therefore most susceptible to the effects of ribosome stacking caused by amino acid shortages (“DISEASE OF CELLULAR PROLIFERATION”). “ Ehlers-Danlos Syndrome”, which belongs to “AORTIC DISEASE”, is caused by “COLLAGEN” gene mutation [44]; the lengthy collagen gene is more affected by ribosome stacking during translation.

**Table 6.** Top 10 diseases in “Jensen DISEASES Curated 2025” category of Enrichr.

| Term | Overlap | P-value | Adjusted P-value |
| --- | --- | --- | --- |
| DISEASE | 447/3737 | $1.27 \times 10^{-12}$ | $5.21 \times 10^{-10}$ |
| DISEASE OF METABOLISM | 112/670 | $4.06 \times 10^{-11}$ | $8.34 \times 10^{-9}$ |
| INHERITED METABOLIC DISORDER | 99/592 | $5.54 \times 10^{-10}$ | $7.60 \times 10^{-8}$ |
| GENETIC DISEASE | 209/1648 | $5.42 \times 10^{-8}$ | $5.57 \times 10^{-6}$ |
| DISEASE OF ANATOMICAL ENTITY | 311/2727 | $1.03 \times 10^{-6}$ | $8.43 \times 10^{-5}$ |
| CANCER | 74/484 | $2.88 \times 10^{-6}$ | $1.98 \times 10^{-4}$ |
| DISEASE OF CELLULAR PROLIFERATION | 74/487 | $3.62 \times 10^{-6}$ | $2.13 \times 10^{-4}$ |
| EHLERS-DANLOS SYNDROME | 9/21 | $3.77 \times 10^{-5}$ | $1.72 \times 10^{-3}$ |
| COLLAGEN DISEASE | 9/21 | $3.77 \times 10^{-5}$ | $1.72 \times 10^{-3}$ |
| AORTIC DISEASE | 8/18 | $7.53 \times 10^{-5}$ | $3.10 \times 10^{-3}$ |

Although we have found more functional enrichment of the selected set of 1,781 genes, it is too much to be interpreted manually. Thus we provide generative AI-based comprehensive analysis as supplementary documents for the readers reference.

### 3.5. *Gene selection: ℓ*_1_ = 3

Next, we consider *u*_3*k*_ in relation to TB. Subsequently, to determine which 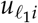 and 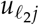 should be considered, we investigate *G*(*ℓ*_1_, *ℓ*_2_, 1, 3) (Fig. 5). Clearly, (*ℓ*_1_, *ℓ*_2_) = (3, 1) is the largest. To confirm the validity of the selection of *u*_1*j*_, we need to see the dependence of *u*_1*j*_ upon *j* (Fig. 3). Although *u*_1*j*_ does not exhibit any *j*-dependence because TB is related to the maintenance of a stable proteome, it is reasonable that *u*_1*j*_ is independent of *j*. Consequently *u*_3*i*_, which shares a large *G* with *u*_1*j*_ and *u*_3*k*_, should be used for gene selection. The left panel of Fig. 6 represents the dependence of the standard deviation of the histogram of 1 − *P*_*i*_ on 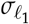. It is obvious that the standard deviation of the histogram of 1 − *P*_*i*_ reaches a minimum at some value of 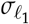. *P*_*i*_ is recomputed using the estimated value of 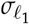. The right panel of Fig. 6 shows the histogram. The large peak on the right side corresponds to the selected genes. *P*-values were corrected using the BH criterion [18], and the 227 genes with adjusted *P*-values less than 0.01 were selected and used for further analysis.

**Figure 5.**
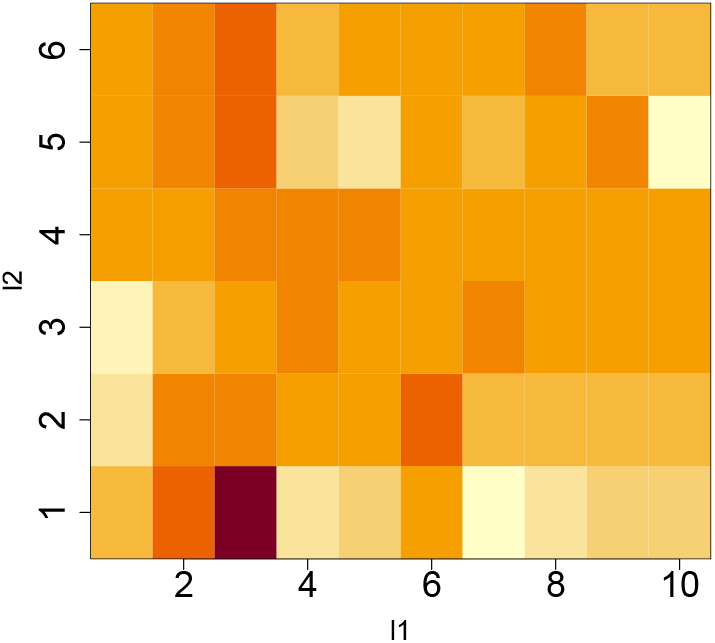
Logarithmic absolute value of *G*(*ℓ*_1_ *ℓ*_2_, 1, 3), log |*G*(*ℓ*_1_, *ℓ*_2_, 1, 3)|. Horizontal:*ℓ*_1_ and vertical: *ℓ*_2_. Darker colors indicate larger values.

**Figure 6.**
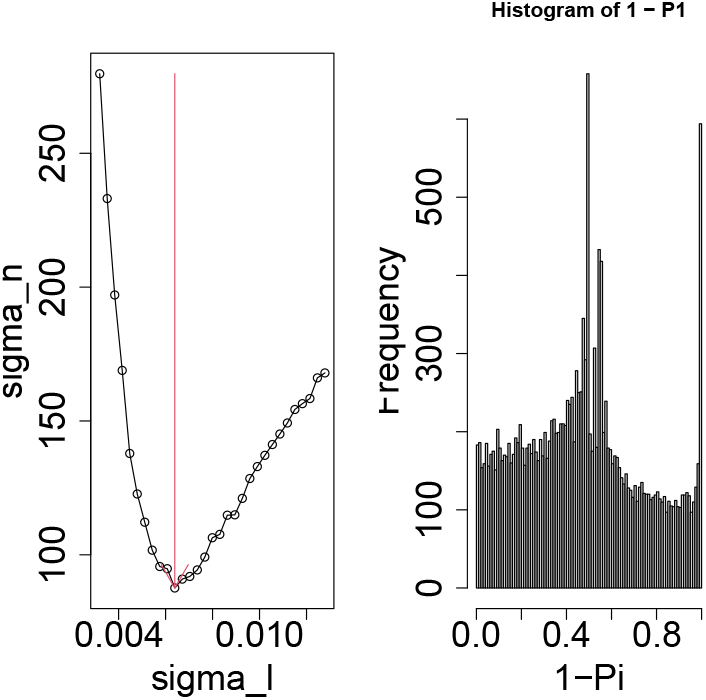
Left: Dependence of the standard deviation (vertical axis) of the histogram of 1 − *P*_*i*_ on 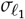 (horizontal axis). Vertical red arrow indicates the estimated 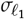. Right: Histogram of 1 − *P*_*i*_ computed using the estimated 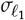.

### 3.6. Validation of translational buffering score

To see whether the selected 227 genes are associated with TB, we defined TB score as

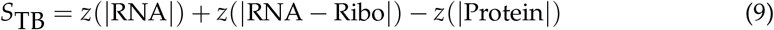

The first term of the right hand was introduced to deny the possibility that genes with missing RNA expression are wrongly selected. We have investigated whether the selected 227 genes are associated with larger *S*TB by applying Wilcoxon rank sum test. This tensor-entry-level comparison showed that the selected 227 genes were associated with larger STB than the remaining genes. (*P* < 2.2 × 10^−16^). Again we did not restrict genes to those with proteome expression, since we could not detect significance only for those with proteome expression. Thus, we can conclude that TD-based unsupervised FE could select the genes associated with TB.

### 3.7. Understanding the biological meaning of the selected genes

To see whether the selected set of 227 genes is biologically meaningful, we have uploaded the selected set of 227 genes to various enrichment analysis servers.

Translational buffering (TB) refers to a regulatory mechanism by which changes in mRNA abundance are offset at the level of translation, ribosome occupancy, or protein output, thereby reducing the direct impact of transcriptional variation on the proteome [45– 49].

Table 7 lists terms related to “Translation” and associated with adjusted *P*-values less than 0.05 in “Reactome Pathways 2024” category of Enrichr. Recent studies [48,49] have reported that the group of genes strongly affected by TB includes many genes related to the “translational machinery” (ribosomes and translational factors), as shown in the Table 7. Especially, it is reasonable that two top pathways are “Eukaryotic Translation Termination” and “Eukaryotic Translation Elongation”, since these two must play critical roles if the selected set of 227 genes are TB-related ones.

**Table 7.**
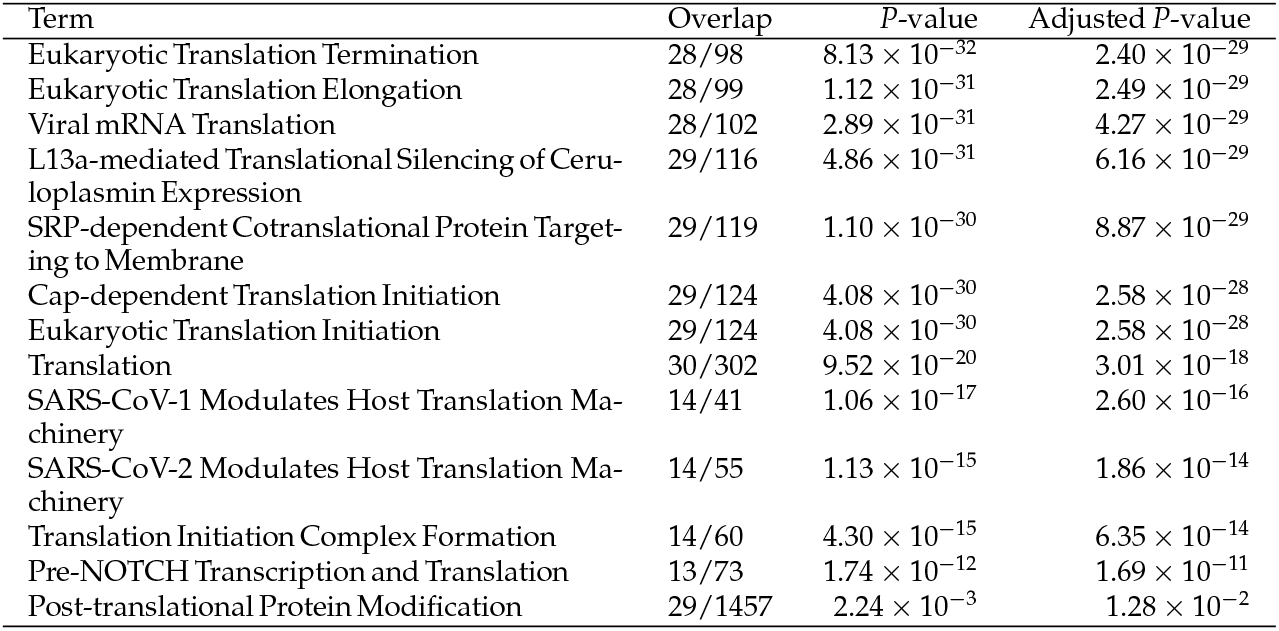
Terms related to “Translation” and associated with adjusted *P*-values less than 0.05 in “Reactome Pathways 2024” category of Enrichr.

Table 8 lists top ranked 10 pathway enrichment results in “KEGG 2026” category in Enrichr. Assuming that TB is occurring, the strong KEGG enrichment of RIBOSOME should be interpreted not merely as enrichment of a cellular component, but as evidence that the selected gene set is connected to the translational layer that can decouple mRNA abundance from protein output.

**Table 8.**
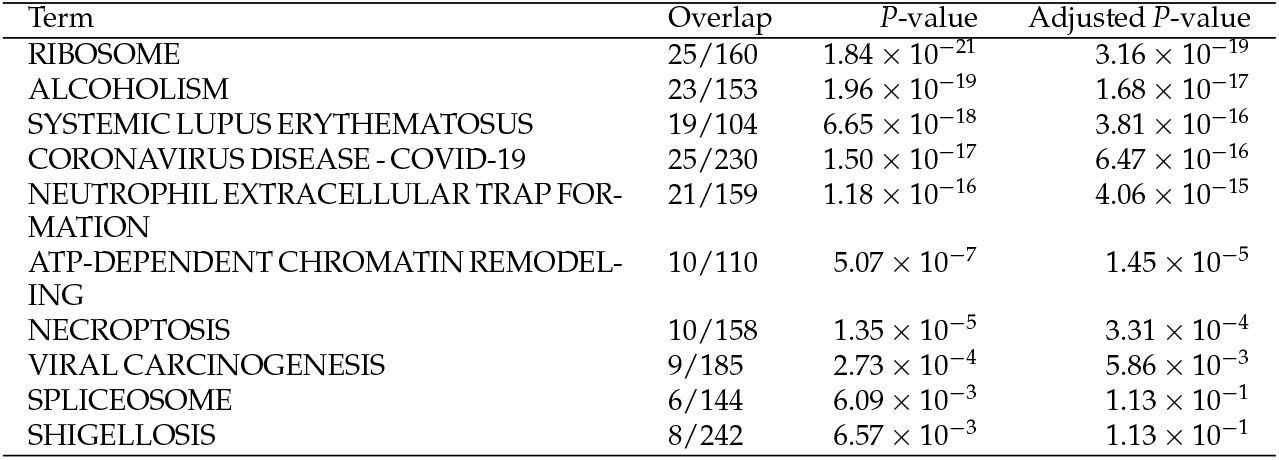
Top ranked 10 pathway enrichment results in “KEGG 2026” category in Enrichr.

In this interpretation, the enrichment of ribosomal genes suggests that the gene set may reflect a compensatory translational response rather than simple downstream protein-level activation. If mRNA levels are broadly altered under disease, stress, or immune activation, cells may adjust ribosome loading, translation initiation, or transcript accessibility so that the resulting protein abundance changes are smaller than the corresponding mRNA changes. This idea is consistent with ribosome-profiling studies, because Ribo-seq directly measures ribosome-protected fragments and therefore provides a more translation-proximal readout than RNA-seq alone [50].

This hypothesis would predict discordance between RNA-seq and Ribo-seq or proteomics: genes contributing to the RIBOSOME term should show large changes at the mRNA level, but smaller or opposite changes in ribosome occupancy, translational efficiency, or protein abundance. Such a pattern has been reported under oxidative stress in yeast, where many genes differentially expressed by RNA-seq were not similarly changed by Ribo-seq, indicating extensive post-transcriptional buffering; notably, translation-related and ribosomal protein genes were among the categories whose mRNA-level changes were compensated at the level of ribosome density [47].

The enrichment of SPLICEOSOME can also be incorporated into the same model, because gene expression is regulated through multiple layers, including transcription, mRNA processing, mRNA stability, translation, and protein stability. In a translational-buffering framework, splicing and RNA-processing changes may represent upstream transcript-level perturbations whose downstream protein-level consequences are partially absorbed by translational control [46,51].

The appearance of infection- and inflammation-related KEGG terms such as COVID-19, viral carcinogenesis, systemic lupus erythematosus, neutrophil extracellular trap formation, and necroptosis can be interpreted as evidence for a stress- or immune-associated context in which translation is actively remodeled. Cellular stress responses commonly involve global repression of protein synthesis together with selective translation of stress-response genes, as seen in the integrated stress response through eIF2*α* phosphorylation [52]. Viral infection is also tightly linked to host translation control; for example, SARS-CoV-2 infection has been reported to reduce host translation and interfere with access of cellular mRNAs to ribosomes [53].

Therefore, under the assumption that TB is operating, the KEGG enrichment profile can be summarized as follows: disease-, immune-, and stress-related transcriptional perturbations may alter mRNA abundance, while ribosome-associated and RNA-processing mechanisms modulate translation so that protein output is buffered. In this view, the top-ranked RIBOSOME term is not merely a pathway annotation, but a mechanistic clue that the selected genes may participate in stabilizing protein abundance despite transcriptomic fluctuation.

Table 9 lists top 10 diseases in “Jensen DISEASES Curated 2025” category of Enrichr. Assuming that translational buffering occurs in the analyzed gene set, the enrichment of Diamond-Blackfan anemia, pure red-cell aplasia, congenital hypoplastic anemia, and related hematopoietic disorders suggests that these genes are linked to ribosomal function, translational control, and erythroid differentiation.

**Table 9.** Top 10 diseases in “Jensen DISEASES Curated 2025” category of Enrichr.

| Term | Overlap | P-value | Adjusted P-value |
| --- | --- | --- | --- |
| PURE RED-CELL APLASIA | 9/25 | $4.65 \times 10^{-12}$ | $3.84 \times 10^{-10}$ |
| DIAMOND-BLACKFAN ANEMIA | 9/25 | $4.65 \times 10^{-12}$ | $3.84 \times 10^{-10}$ |
| CONGENITAL HYPOPLASTIC ANEMIA | 9/47 | $2.50 \times 10^{-9}$ | $1.03 \times 10^{-7}$ |
| APLASTIC ANEMIA | 9/47 | $2.50 \times 10^{-9}$ | $1.03 \times 10^{-7}$ |
| NORMOCYTIC ANEMIA | 10/87 | $5.43 \times 10^{-8}$ | $1.79 \times 10^{-6}$ |
| ANEMIA | 10/94 | $1.14 \times 10^{-7}$ | $3.15 \times 10^{-6}$ |
| HEMATOPOIETIC SYSTEM DISEASE | 11/170 | $4.02 \times 10^{-6}$ | $9.46 \times 10^{-5}$ |
| PHYSICAL DISORDER | 13/438 | $1.56 \times 10^{-3}$ | $3.22 \times 10^{-2}$ |
| MYOPIA | 2/7 | $2.59 \times 10^{-3}$ | $4.76 \times 10^{-2}$ |
| REFRACTIVE ERROR | 2/11 | $6.59 \times 10^{-3}$ | $1.09 \times 10^{-1}$ |

In this context, the strong enrichment of Diamond-Blackfan anemia is particularly informative, because Diamond-Blackfan anemia is a prototypical ribosomopathy caused in many cases by haploinsufficiency of ribosomal protein genes and defects in ribosome biogenesis [54–56].

Thus, the disease enrichment pattern can be interpreted as a disease-level manifestation of disturbed translational homeostasis. If transcript abundance changes are normally buffered at the translational level, then genes involved in ribosome function, ribosome biogenesis, and translational efficiency would be expected to appear prominently among buffered genes. Failure, overload, or dysregulation of this buffering system could preferentially affect erythroid progenitors, because erythropoiesis is highly dependent on coordinated ribosome production and protein synthesis. This is consistent with studies showing that ribosomal protein haploinsufficiency selectively activates stress responses in human erythroid progenitors and impairs erythroid development [57,58].

Moreover, ribosomal protein deficiency does not necessarily reduce translation uniformly across all mRNAs. Instead, reduced ribosome levels can selectively impair translation of specific transcripts important for hematopoietic lineage commitment, including erythroid regulators such as GATA1. Altered translation of GATA1 has been proposed as a key mechanism linking ribosomal protein haploinsufficiency to the erythroid defect in Diamond-Blackfan anemia [59,60].

Therefore, under the assumption of TB, the enrichment of anemia-related and ribosomopathy-related terms does not simply indicate a generic association with blood disorders. Rather, it suggests that the analyzed gene set may represent a translationally buffered network in which perturbations of ribosome function, translational efficiency, and erythroid differentiation are mechanistically connected. The enrichment of Diamond-Blackfan anemia provides the most direct link, whereas broader terms such as aplastic anemia, normocytic anemia, anemia, and hematopoietic system disease may reflect down-stream or more general consequences of impaired translational homeostasis in hematopoietic cells.

In conclusion, it is quite reasonable to assume that the selected set of 227 genes is related to TB.

## 4. Discussion

We applied TD-based unsupervised FE to tri-omics comprising the transcriptome, translatome, and proteome. The 1,781 and 227 genes that are supposed to be associated with ribosome stacking and TB, respectively, were enriched in reasonable biological terms. In spite of that, there might be some concerns about pre-processing. We address these possible methodological concerns in advance.

At first, we intentionally replicated one of two samples of proteome since there were only two samples whereas there are three samples for other experimental condition. To see whether this duplication skewed the results, we replicate another of two samples. We have found that *u*_3*i*_ and *u*_5*i*_ when another of two samples is replicated correspond to *u*_3*i*_ and *u*_6*i*_ (Fig. 7). It is obvious that *u*_3*i*_s are identical with each other between distinct ways of replication. Although not *u*_6*i*_ but *u*_5*i*_ is similar to *u*_6*i*_ when the first way of replication, whether the same 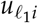 correspond to each other does not matter since we can freely select the suitable one. Since *u*_6*i*_ a bit differs from *u*_5*i*_, we further compared the selected genes between two ways of replication (Table 10). Since the more than 90 % of the selected genes are common between two ways of replication, it is obvious that which one of samples is replicated does not matter.

**Table 10.** Confusion matrix between the selected genes bwteen two ways of replication. Rows: the first one, columns: the second one

| | | Adjusted P-values using $u_{5i}$ | |
| --- | --- | --- | --- |
| | | $\geq 0.01$ | $\leq 0.01$ |
| Adjusted P-values using $u_{6i}$ | $\geq 0.01$ | 16296 | 98 |
| | $\leq 0.01$ | 153 | 1628 |

**Figure 7.**
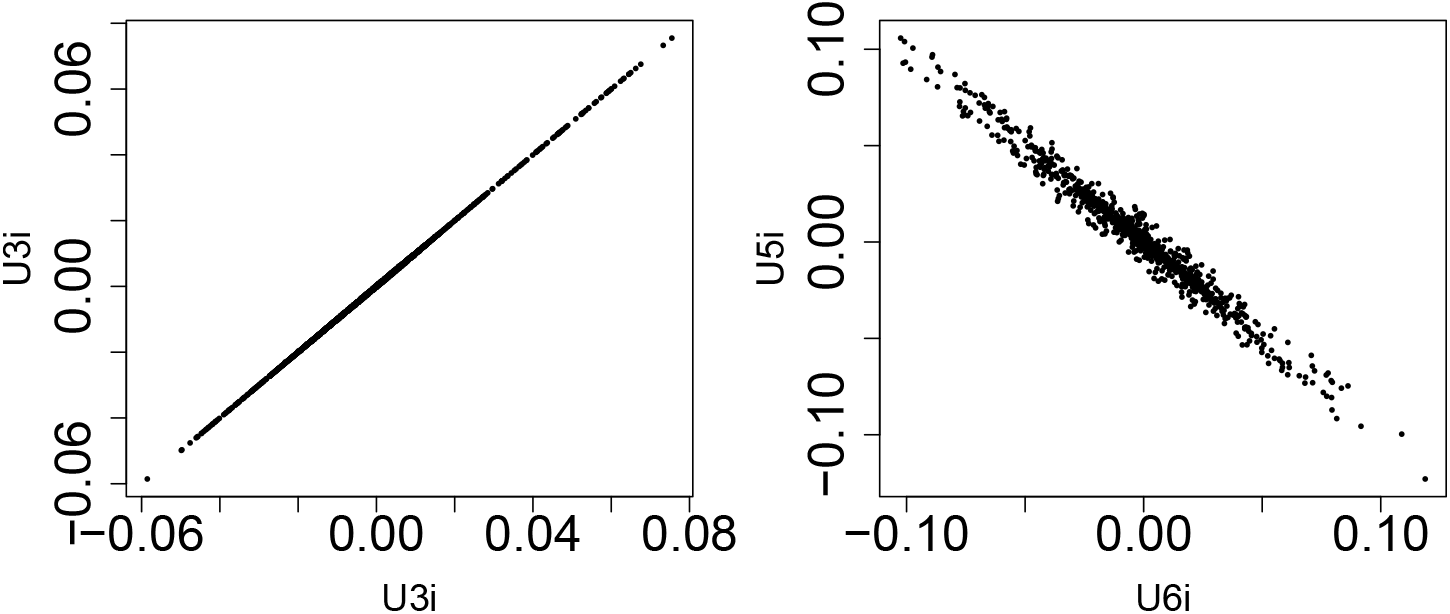
Comparison of singular value vectors between replications of different samples. Left: *u*_3*i*_ when the first one is replicated (horizontal) and that when the second one is replicated (vertical). Left: *u*_6*i*_ when the first one is replicated (horizontal) and *u*_5*i*_ when the second one is replicated (vertical).

Yet another concern is to fill zero to missing values in proteome. To see the effect of this pre-process, we employed the alternative strategy at the very last stage (i.e., just before applying TD).

1. Filling zero to missing proteome values
2. Follow the pre-process as suggested in the paper.
3. At the very last stage, *i*s whose proteome is missing is replaced with not missing values as follows.
4. Compute Euclidean distance between the corresponding gene expression or translatome (i.e., for *i*th one) and all other gene expression or translatome (i.e., for all *i′* not equal to *i*).
5. Find *i′* whose gene expression or translatome is the closest to that associated with missing proteome, *i*.
6. Substitute the corresponding proteome (i.e., for the identified *i′*the one) to the missing values (i.e. *i*th one) of proteome.

One should recognize that it is extensive alteration, since the number of substitution is larger than half of proteome (because the number of proteome provided is 7,004, which is much less than *N*(=18,175), the number of genes/translatome) and missing proteome is supposed to be zero (i.e., not detected at all). In spite of the extensive alteration, there are still the corresponding 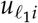 s (Fig. 8). One might wonder whether it can happen since more than half of proteome values changed. This can be understood as follows. In principle, not all *i*s contribute to a specific 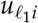. In contrast, eq. (5) assumes that only limited part of genes contribute to individual 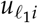 and can be therefore selected. Table 11 shows the confusion matrix between genes selected and genes whose proteome is missing. Regardless to zero filling or alternative substitution, genes whose proteome is missing is rarely selected. This suggests that how to fill the missing values of proteome does not affect the results in this paper so much.

**Table 11.** Confusion matrix between the selected genes between zero filling or alternative substitution and genes whose proteome is missing. Rows: genes whose proteome is missing, columns: the selected genes,

| | Adjusted P-values using $u_{6i}$ | | Adjusted P-values using $u_{5i}$ | |
| --- | --- | --- | --- | --- |
|  | zero filling |  | alternative substitution |  |
| | $\geq 0.01$ | $\leq 0.01$ | $\geq 0.01$ | $\leq 0.01$ |
| Proteome not missing | 4846 | 1781 | 5926 | 701 |
| Proteome missing | 11548 | 0 | 11501 | 47 |

**Figure 8.**
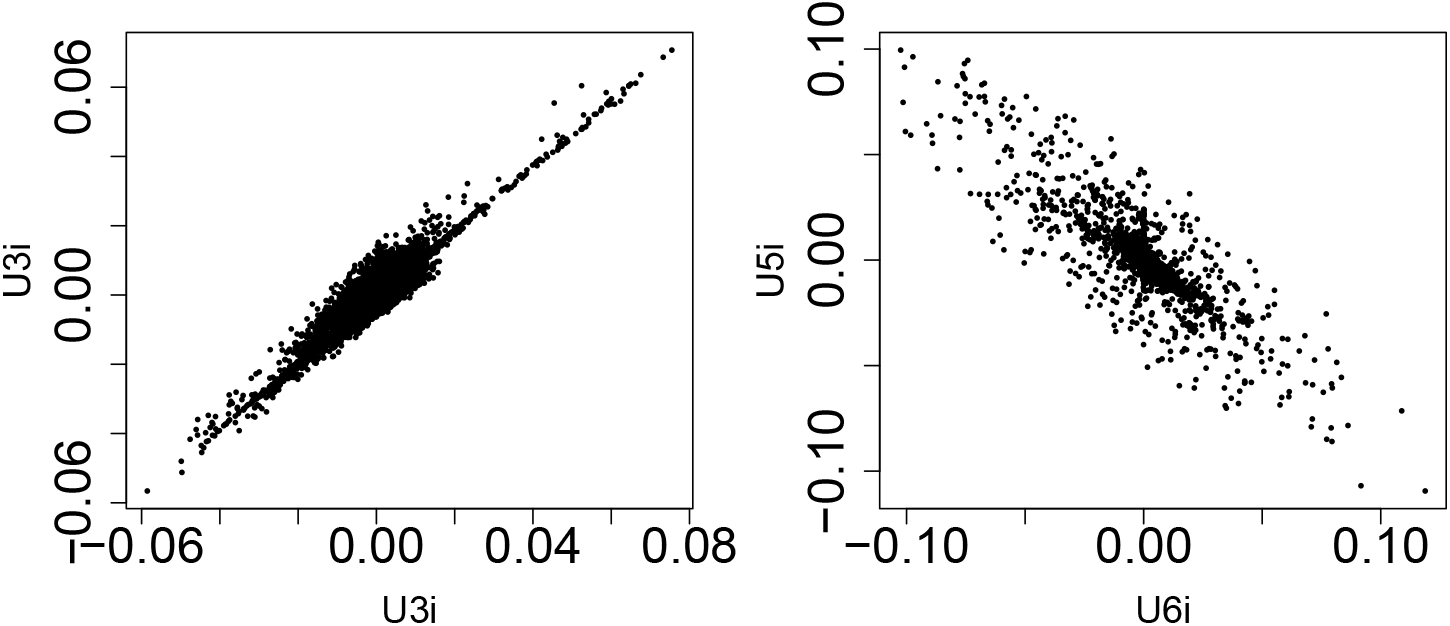
Comparison of singular value vectors between zero filling and the alternative substitution. Left: *u*_3*i*_ when for zero filling(horizontal) and that for the alternative substitution (vertical). Right: *u*_6*i*_ for zero filling (horizontal) and *u*_5*i*_ for the alternative substitution (vertical).

A natural question is whether existing multi-omics integration methods can recover similar components.

For fair comparison, MOFA+ and mixOmics were applied to the same preprocessed data used immediately before TD-based unsupervised FE, as described in Section 2.6. Under the tested settings and the evaluation criteria used here, mixOmics did not recover components clearly corresponding to ribosome stacking or TB. Therefore, TD-based unsupervised FE yielded more directly interpretable components corresponding to ribosome stacking and TB than MOFA+ or mixOmics (Fig. 9). For clarification, numerical values corresponding to Fig. 9 are also provided in Table S1.

**Figure 9.**
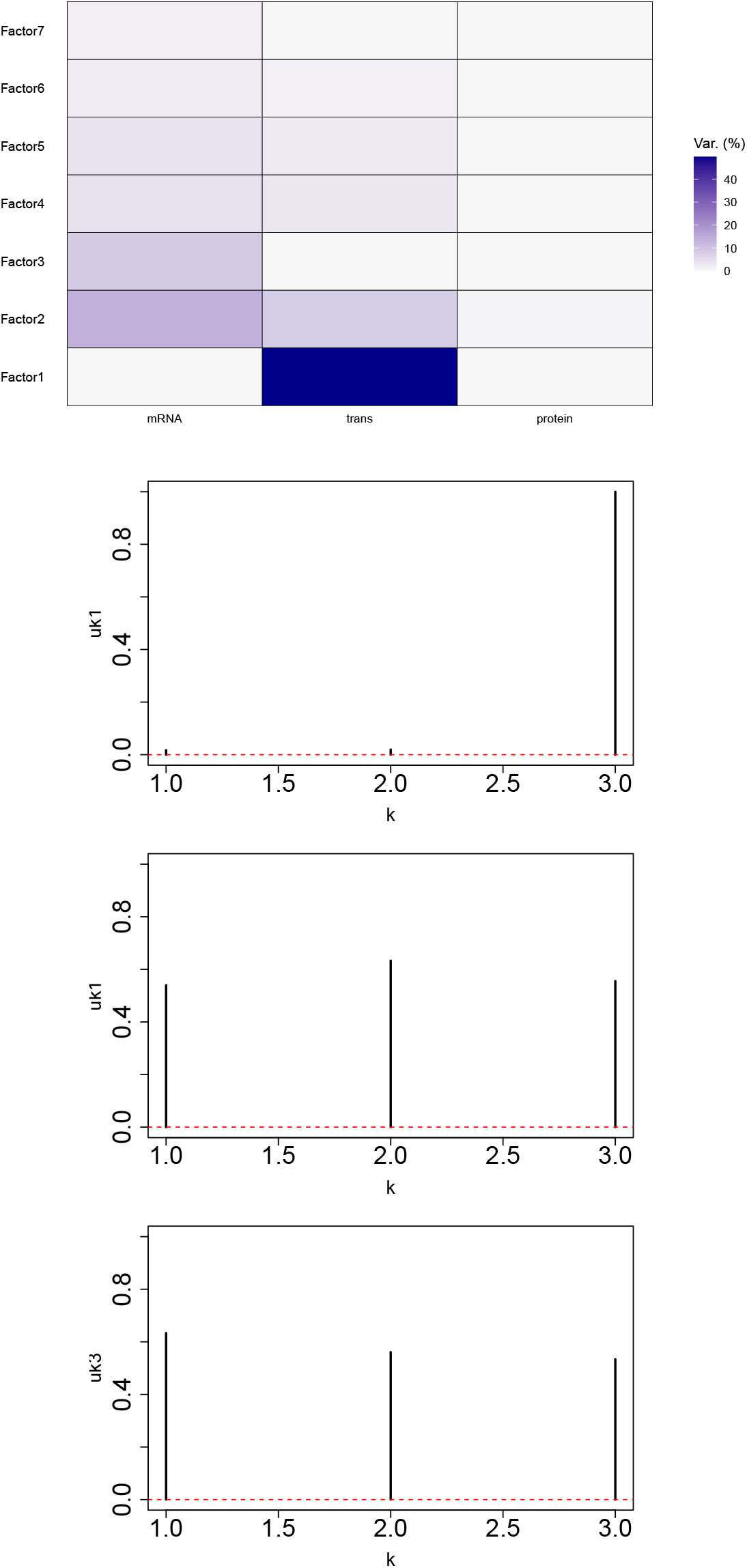
The performances by SOTA. Top row: weight (row: components, column:tri-omics) matrix by MOFA+. Second to fourth rows: the first three components by mixOmics. *k* = 1: transcriptome, *k* = 2: translatome, *k* = 3: proteome.

This suggests how difficult to get components consistent with ribosome stacking or TB using the present datasets, whereas TD identified interpretable omics-mode vectors corresponding to these patterns (Fig. 1).

## 5. Conclusions

In this study, we developed and applied a four-dimensional tensor decomposition-based unsupervised feature extraction method to integrate transcriptome, translatome, and proteome profiles obtained under branched-chain amino acid starvation. By representing genes, experimental conditions, replicates, and omics layers as a single tensor, the proposed method enabled the direct extraction of components reflecting relationships among mRNA abundance, ribosome occupancy, and protein abundance.

The analysis identified two biologically interpretable components. One component was consistent with ribosome stacking, because it represented increased transcriptome and translatome signals accompanied by decreased proteome signals. Gene selection based on this component identified 1,781 genes enriched for translation-related processes, post-translational protein modification, transcriptional regulation, cell cycle suppression, endoplasmic reticulum protein processing, ubiquitin-mediated proteolysis, and stress-response pathways. These enrichments suggest that the selected genes reflect cellular responses to impaired translation caused by amino acid starvation and ribosomal congestion.

The second component was consistent with translational buffering, because it represented a pattern in which proteome abundance remained relatively stable despite changes in transcriptome and translatome profiles. Gene selection based on this component identified 227 genes enriched for ribosome-related pathways, eukaryotic translation elongation and termination, spliceosome, immune- and stress-associated pathways, and anemia- or ribosomopathy-related disease terms. These results suggest that the selected genes are involved in regulatory mechanisms that buffer protein abundance against transcriptomic perturbation.

Robustness analyses further supported the reliability of the proposed framework. The major singular value vectors and selected genes were largely preserved when a different proteome replicate was duplicated or when missing proteome values were handled using an alternative substitution strategy. Moreover, comparison with MOFA+ and mixOmics indicated that these widely used multi-omics integration methods did not extract components as clearly consistent with ribosome stacking or translational buffering as those obtained by tensor decomposition.

Taken together, these findings indicate that tensor decomposition-based unsupervised feature extraction can effectively integrate tri-omics profiles and identify gene clusters with coherent biological and functional interpretations. The proposed approach provides a useful data-driven framework for investigating translational regulation, ribosome stacking, and translational buffering in multi-layer omics datasets. Future studies applying this method to additional biological conditions and experimentally validating the selected genes will further clarify its utility for understanding post-transcriptional and translational regulation.

## Supplementary Materials

The following supporting information can be downloaded at: https://www.mdpi.com/article/10.3390/biology1010000/s1,

Supplementary file 1 (Excel): List of genes selected by *ℓ*_1_ = 3 and 6, and the associated enrichment clustering provided by DAVID.

Supplementary file 2 (PDF): Generative AI based summary of enrichment analysis only for the set of the selected 1,781 genes.

Supplementary file 3 (PDF): Table S1

## Author Contributions

Y. H. T. planned the study and performed the analyses. Y.-H.T. and T.T. evaluated the results and wrote and reviewed the manuscript. Conceptualization, data curation, and analysis were performed by Y. H. T. All the authors have read and agreed to the published version of the manuscript.

## Institutional Review Board Statement

NA

## Informed Consent Statement

NA

## Data Availability Statement

Transcriptome and Ribo-seq data are available through GEO accession numbers GSE291652 and GSE291653, respectively. Proteome data are available through PRIDE accession number PXD067949. Sample code is in https://github.com/tagtag/TDbasedUFE_Ribo-seq.

## Disclaimer/Publisher’s Note

The statements, opinions and data contained in all publications are solely those of the individual author(s) and contributor(s) and not of MDPI and/or the editor(s). MDPI and/or the editor(s) disclaim responsibility for any injury to people or property resulting from any ideas, methods, instructions or products referred to in the content.

## References

1. Liu, Y.; Beyer, A.; Aebersold, R. On the Dependency of Cellular Protein Levels on mRNA Abundance. Cell 2016, 165, 535–550. 10.1016/j.cell.2016.03.014.

2. Brito Querido, J.; Díaz-López, I.; Ramakrishnan, V. The molecular basis of translation initiation and its regulation in eukaryotes. Nature Reviews Molecular Cell Biology 2024, 25, 168–186. 10.1038/s41580-023-00624-9.

3. Joazeiro, C.A.P. Mechanisms and functions of ribosome-associated protein quality control. Nature Reviews Molecular Cell Biology 2019, 20, 368–383. 10.1038/s41580-019-0118-2.

4. Limbu, M.S.; Xiong, T.; Wang, S. A review of Ribosome profiling and tools used in Ribo-seq data analysis. Computational and Structural Biotechnology Journal 2024, 23, 1912–1918. 10.1016/j.csbj.2024.04.051.

5. Schwanhäusser, B.; Busse, D.; Li, N.; Dittmar, G.; Schuchhardt, J.; Wolf, J.; Chen, W.; Selbach, M. Global quantification of mammalian gene expression control. Nature 2011, 473, 337–342. 10.1038/nature10098.

6. Baysoy, A.; Bai, Z.; Satija, R.; Fan, R. The technological landscape and applications of single-cell multi-omics. Nature Reviews Molecular Cell Biology 2023, 24, 695–713. 10.1038/s41580-023-00615-w.

7. Jovanovic, M.; Rooney, M.S.; Mertins, P.; Przybylski, D.; Chevrier, N.; Satija, R.; Rodriguez, E.H.; Fields, A.P.; Schwartz, S.; Raychowdhury, R.; et al. Dynamic profiling of the protein life cycle in response to pathogens. Science 2015, 347, 1259038, [https://www.science.org/doi/pdf/10.1126/science.1259038]. 10.1126/science.1259038.

8. Rao, S.; Le, A.Y.; Persyn, L.; Cenik, C. Translational buffering tunes gene expression in mice and humans. Genome Biology 2026, 27, 113. 10.1186/s13059-026-04010-4.

9. Battle, A.; Khan, Z.; Wang, S.H.; Mitrano, A.; Ford, M.J.; Pritchard, J.K.; Gilad, Y. Impact of regulatory variation from RNA to protein. Science 2015, 347, 664–667, [https://www.science.org/doi/pdf/10.1126/science.1260793]. 10.1126/science.1260793.

10. Cenik, C.; Cenik, E.S.; Byeon, G.W.; Grubert, F.; Candille, S.I.; Spacek, D.; Alsallakh, B.; Tilgner, H.; Araya, C.L.; Tang, H.; et al. Integrative analysis of RNA, translation, and protein levels reveals distinct regulatory variation across humans. Genome Research 2015, 25, 1610–1621, [http://genome.cshlp.org/content/25/11/1610.full.pdf+html]. 10.1101/gr.193342.115.

11. Blevins, W.R.; Tavella, T.; Moro, S.G.; Blasco-Moreno, B.; Closa-Mosquera, A.; Díez, J.; Carey, L.B.; M, M.A. Extensive post-transcriptional buffering of gene expression in the response to severe oxidative stress in baker’s yeast. Scientific Reports 2019, 9, 11005. 10.1038/s41598-019-47424-w.

12. Carlyle, B.C.; Kitchen, R.R.; Zhang, J.; Wilson, R.S.; Lam, T.T.; Rozowsky, J.S.; Williams, K.R.; Sestan, N.; Gerstein, M.B.; Nairn, A.C. Isoform-Level Interpretation of High-Throughput Proteomics Data Enabled by Deep Integration with RNA-seq. Journal of Proteome Research 2018, 17, 3431–3444. 10.1021/acs.jproteome.8b00310.

13. Liu, T.Y.; Huang, H.H.; Wheeler, D.; Xu, Y.; Wells, J.A.; Song, Y.S.; Wiita, A.P. Time-Resolved Proteomics Extends Ribosome Profiling-Based Measurements of Protein Synthesis Dynamics. Cell Systems 2017, 4, 636–644.e9. 10.1016/j.cels.2017.05.001.

14. Cuevas, R.; et al. Most non-canonical proteins uniquely populate the proteome or immunopeptidome. Cell Reports 2021,34, 108815. 10.1016/j.celrep.2021.108815.

15. Zhu, W.; Xu, J.; Chen, S.; Chen, J.; Liang, Y.; Zhang, C.; Li, Q.; Lai, J.; Li, L. Large-scale translatome profiling annotates the functional genome and reveals the key role of genic 3’ untranslated regions in translatomic variation in plants. Plant Communications 2021, 2, 100181. 10.1016/j.xplc.2021.100181.

16. Worpenberg, L.; Gobet, C.; Naef, F. Codon-specific ribosome stalling reshapes translational dynamics during branched-chain amino acid starvation. Genome Biology 2025, 26, 315. 10.1186/s13059-025-03800-6.

17. Statoulla, E.; Zafeiri, M.; Chalkiadaki, K.; Voudouri, G.; Gkika, K.S.; Papazoglou, C.; Durcan, T.M.; Khoutorsky, A.; Jafarnejad, S.M.; Maguire, S.; et al. SNCA triplication disrupts proteostasis and extracellular architecture prior to neurodegeneration in human midbrain organoids. npj Parkinson’s Disease 2026. 10.1038/s41531-026-01292-0.

18. Taguchi, Y.h. Unsupervised Feature Extraction Applied to Bioinformatics: A PCA Based and TD Based Approach, 2 ed.; Unsupervised and Semi-Supervised Learning, Springer International Publishing: Switzland, 2024. 10.1007/978-3-031-60982-4.

19. Wang, X.; Wang, C.; Ji, B.; Wang, J.; Zheng, M.; Song, L.; Peng, S.; Shang, X. Multimodal pre-training models of molecular representation for drug discovery. National Science Review 2026, 13, waf495, [https://academic.oup.com/nsr/article-pdf/13/1/nwaf495/65269418/nwaf495.pdf]. 10.1093/nsr/nwaf495.

20. Wei, M.M.; Wang, L.; Zhao, B.W.; Su, X.R.; You, Z.H.; Huang, D.S. Integrating Transformer and Graph Attention Network for circRNA-miRNA Interaction Prediction. IEEE Journal of Biomedical and Health Informatics 2025, 29, 6105–6113. 10.1109/JBHI.2025.3561197.

21. Wei, M.; Wang, L.; Su, X.; Zhao, B.; You, Z. Multi-hop graph structural modeling for cancer-related circRNA-miRNA interaction prediction. Pattern Recognition 2026, 170, 112078. 10.1016/j.patcog.2025.112078.

22. Ji, B.; Hu, T.; Wang, J.; Liu, M.; Xu, L.; Zhang, Q.; Zhang, Y.; Qiao, L.; Zhang, Y.; Peng, S.; et al. CAPTAIN: a multimodal foundation model pretrained on co-assayed single-cell RNA and protein. Nature Communications 2026. 10.1038/s41467-026-72882-y.

23. Cui, Y.; Peng, C.; Xia, Z.; Yang, C.; Guo, Y. A survey of sequence-to-graph mapping algorithms in the pangenome era. Genome Biology 2025, 26, 138. 10.1186/s13059-025-03606-6.

24. Perez-Riverol, Y.; Bai, M.; Da Veiga Leprevost, F.; et al. The PRIDE database resources in 2025: 20 years of proteomics data sharing. Nucleic Acids Research 2025, 53, D1528–D1539. 10.1093/nar/gkae1026.

25. Durinck, S.; Spellman, P.T.; Birney, E.; Huber, W. Mapping identifiers for the integration of genomic datasets with the R/Bioconductor package biomaRt. Nature Protocols 2009, 4, 1184–1191. 10.1038/nprot.2009.97.

26. Sherman, B.T.; Hao, M.; Qiu, J.; Jiao, X.; Baseler, M.W.; Lane, H.C.; Imamichi, T.; Chang, W. DAVID: a web server for functional enrichment analysis and functional annotation of gene lists (2021 update). Nucleic Acids Research 2022, 50, W216–W221. 10.1093/nar/gkac194.

27. Xie, Z.; Bailey, A.; Kuleshov, M.V.; Clarke, D.J.B.; Evangelista, J.E.; Jenkins, S.L.; Lachmann, A.; Wojciechowicz, M.L.; Kropiwnicki, E.; Jagodnik, K.M.; et al. Gene Set Knowledge Discovery with Enrichr. Current Protocols 2021, 1, e90, [https://currentprotocols.onlinelibrary.wiley.com/doi/pdf/10.1002/cpz1.90]. 10.1002/cpz1.90.

28. Argelaguet, R.; Arnol, D.; Bredikhin, D.; Deloro, Y.; Velten, B.; Marioni, J.C.; Stegle, O. MOFA+: a statistical framework for comprehensive integration of multi-modal single-cell data. Genome Biology 2020, 21, 111. 10.1186/s13059-020-02015-1.

29. Rohart, F.; Gautier, B.; Singh, A.; Lê Cao, K.A. mixOmics: An R package for ‘omics feature selection and multiple data integration. PLOS Computational Biology 2017, 13, 1–19. 10.1371/journal.pcbi.1005752.

30. De, S.; Mühlemann, O. A comprehensive coverage insurance for cells: revealing links between ribosome collisions, stress responses and mRNA surveillance. RNA Biology 2022, 19, 609–621, [https://doi.org/10.1080/15476286.2022.2065116]. PMID: 35491909, 10.1080/15476286.2022.2065116.

31. Bhawe, K.; Roy, D. Interplay between NRF1, E2F4 and MYC transcription factors regulating common target genes contributes to cancer development and progression. Cellular Oncology 2018, 41, 465–484. 10.1007/s13402-018-0395-3.

32. Vind, A.C.; Genzor, A.V.; Bekker-Jensen, S. Ribosomal stress-surveillance: three pathways is a magic number. Nucleic Acids Research 2020, 48, 10648–10661, [https://academic.oup.com/nar/article-pdf/48/19/10648/34133758/gkaa757.pdf]. 10.1093/nar/gkaa757.

33. Louder, R.K.; He, Y.; López-Blanco, J.R.; Fang, J.; Chacón, P.; Nogales, E. Structure of promoter-bound TFIID and model of human pre-initiation complex assembly. Nature 2016, 531, 604–609. 10.1038/nature17394.

34. Grzenda, A.; Lomberk, G.; Zhang, J.S.; Urrutia, R. Sin3: Master scaffold and transcriptional corepressor. Biochimica et Biophysica Acta (BBA) - Gene Regulatory Mechanisms 2009, 1789, 443–450. 10.1016/j.bbagrm.2009.05.007.

35. Weintraub, A.S.; Li, C.H.; Zamudio, A.V.; Sigova, A.A.; Hannett, N.M.; Day, D.S.; Abraham, B.J.; Cohen, M.A.; Nabet, B.; Buckley, D.L.; et al. YY1 Is a Structural Regulator of Enhancer-Promoter Loops. Cell 2017, 171, 1573–1588.e28. 10.1016/j.cell.2017.11.008.

36. Jobava, R.; Mao, Y.; Guan, B.J.; Hu, D.; Krokowski, D.; Chen, C.W.; Shu, X.E.; Chukwurah, E.; Wu, J.; Gao, Z.; et al. Adaptive translational pausing is a hallmark of the cellular response to severe environmental stress. Molecular Cell 2021, 81, 4191–4208.e8. 10.1016/j.molcel.2021.09.029.

37. Gandin, V.; Topisirovic, I. Co-translational mechanisms of quality control of newly synthesized polypeptides. Translation 2014, 2, e28109. [https://doi.org/10.4161/trla.28109]. PMID: 26779401, 10.4161/trla.28109.

38. Kikushige, Y.; Miyamoto, T.; Kochi, Y.; Semba, Y.; Ohishi, M.; Irifune, H.; Hatakeyama, K.; Kunisaki, Y.; Sugio, T.; Sakoda, T.; et al. Human acute leukemia uses branched-chain amino acid catabolism to maintain stemness through regulating PRC2 function. Blood Advances 2023, 7, 3592–3603, [https://ashpublications.org/bloodadvances/article-pdf/7/14/3592/2065310/blooda_adv-2022-008242-main.pdf]. 10.1182/bloodadvances.2022008242.

39. Yu, F.X.; Zhao, B.; Guan, K.L. Hippo Pathway in Organ Size Control, Tissue Homeostasis, and Cancer. Cell 2015, 163, 811–828. 10.1016/j.cell.2015.10.044.

40. Vanweert, F.; Schrauwen, P.; Phielix, E. Role of branched-chain amino acid metabolism in the pathogenesis of obesity and type 2 diabetes-related metabolic disturbances BCAA metabolism in type 2 diabetes. Nutrition & Diabetes 2022, 12, 35. 10.1038/s41387-022-00213-3.

41. Ishimura, R.; Nagy, G.; Dotu, I.; Zhou, H.; Yang, X.L.; Schimmel, P.; Senju, S.; Nishimura, Y.; Chuang, J.H.; Ackerman,S.L. Ribosome stalling induced by mutation of a CNS-specific tRNA causes neurodegeneration. Science 2014, 345, 455–459, [https://www.science.org/doi/pdf/10.1126/science.1249749]. 10.1126/science.1249749.

42. Ni, C.; Buszczak, M. Ribosome biogenesis and function in development and disease. Development 2023, 150, dev201187. [https://journals.biologists.com/dev/article-pdf/150/5/dev201187/2639866/dev201187.pdf]. 10.1242/dev.201187.

43. Tönjes, M.; Barbus, S.; Park, Y.J.; Wang, W.; Schlotter, M.; Lindroth, A.M.; Pleier, S.V.; Bai, A.H.C.; Karra, D.; Piro, R.M.; et al. BCAT1 promotes cell proliferation through amino acid catabolism in gliomas carrying wild-type IDH1. Nature Medicine 2013, 19, 901–908. 10.1038/nm.3217.

44. Malfait, F.; Francomano, C.; Byers, P.; Belmont, J.; Berglund, B.; Black, J.; Bloom, L.; Bowen, J.M.; Brady, A.F.; Burrows, N.P.; et al. The 2017 international classification of the Ehlers–Danlos syndromes. American Journal of Medical Genetics Part C: Seminars in Medical Genetics 2017, 175, 8–26, [https://onlinelibrary.wiley.com/doi/pdf/10.1002/ajmg.c.31552]. 10.1002/ajmg.c.31552.

45. McManus, C.J.; May, G.E.; Spealman, P.; Shteyman, A. Ribosome profiling reveals post-transcriptional buffering of divergent gene expression in yeast. Genome Research 2014, 24, 422–430, [http://genome.cshlp.org/content/genome/24/3/422.full.pdf]. 10.1101/gr.164996.113.

46. Liu, Y.; Beyer, A.; Aebersold, R. On the Dependency of Cellular Protein Levels on mRNA Abundance. Cell 2016, 165, 535–550. 10.1016/j.cell.2016.03.014.

47. Blevins, W.R.; Tavella, T.; Moro, S.G.; Blasco-Moreno, B.; Closa-Mosquera, A.; Díez, J.; Carey, L.B.; M, M.A. Extensive post-transcriptional buffering of gene expression in the response to severe oxidative stress in baker’s yeast. Scientific Reports 2019, 9, 11005. 10.1038/s41598-019-47424-w.

48. Kusnadi, E.P.; Timpone, C.; Topisirovic, I.; Larsson, O.; Furic, L. Regulation of gene expression via translational buffering. Biochimica et Biophysica Acta (BBA) - Molecular Cell Research 2022, 1869, 119140. 10.1016/j.bbamcr.2021.119140.

49. Rao, S.; Le, A.Y.; Persyn, L.; Cenik, C. Translational buffering tunes gene expression in mice and humans. Genome Biology 2026,27, 113. 10.1186/s13059-026-04010-4.

50. Ingolia, N.T.; Ghaemmaghami, S.; Newman, J.R.S.; Weissman, J.S. Genome-Wide Analysis in Vivo of Translation with Nucleotide Resolution Using Ribosome Profiling. Science 2009, 324, 218–223, [https://www.science.org/doi/pdf/10.1126/science.1168978]. 10.1126/science.1168978.

51. Oertlin, C.; Lorent, J.; Murie, C.; Furic, L.; Topisirovic, I.; Larsson, O. Generally applicable transcriptome-wide analysis of translation using anota2seq. Nucleic Acids Research 2019, 47, e70.–e70, [https://academic.oup.com/nar/articlepdf/47/12/e70/28917189/gkz223.pdf]. 10.1093/nar/gkz223.

52. Pakos-Zebrucka, K.; Koryga, I.; Mnich, K.; Ljujic, M.; Samali, A.; Gorman, A.M. The integrated stress response. The EMBO Reports 2016, 17, 1374–1395. 10.15252/embr.201642195.

53. Finkel, Y.; Gluck, A.; Nachshon, A.; Winkler, R.; Fisher, T.; Rozman, B.; Mizrahi, O.; Lubelsky, Y.; Zuckerman, B.; Slobodin, B.; et al. SARS-CoV-2 uses a multipronged strategy to impede host protein synthesis. Nature 2021, 594, 240–245. 10.1038/s41586-021-03610-3.

54. Narla, A.; Ebert, B.L. Ribosomopathies: human disorders of ribosome dysfunction. Blood 2010, 115, 3196–3205, [https://ashpublications.org/blood/article-pdf/115/16/3196/1325052/zh801610003196.pdf]. 10.1182/blood-2009-10-178129.

55. Da Costa, L.; Leblanc, T.; Mohandas, N. Diamond-Blackfan anemia. Blood 2020, 136, 1262–1273, [https://ashpublications.org/blood/pdf/136/11/1262/1757318/bloodbld2019000947c.pdf]. 10.1182/blood.2019000947.

56. Da Costa, L.; Mohandas, N.; David-NGuyen, L.; Platon, J.; Marie, I.; O’Donohue, M.F.; Leblanc, T.; Gleizes, P.E. Diamond-Blackfan anemia, the archetype of ribosomopathy: How distinct is it from the other constitutional ribosomopathies? Blood Cells, Molecules, and Diseases 2024, 106, 102838. 10.1016/j.bcmd.2024.102838.

57. Dutt, S.; Narla, A.; Lin, K.; Mullally, A.; Abayasekara, N.; Megerdichian, C.; Wilson, F.H.; Currie, T.; Khanna-Gupta, A.; Berliner, N.; et al. Haploinsufficiency for ribosomal protein genes causes selective activation of p53 in human erythroid progenitor cells. Blood 2011, 117, 2567–2576, [https://ashpublications.org/blood/article-pdf/117/9/2567/1315179/zh800911002567.pdf]. 10.1182/blood-2010-07-295238.

58. Narla, A.; Ebert, B.L. Ribosomopathies: human disorders of ribosome dysfunction. Blood 2010, 115, 3196–3205, [https://ashpublications.org/blood/article-pdf/115/16/3196/1325052/zh801610003196.pdf]. 10.1182/blood-2009-10-178129.

59. Ludwig, L.S.; Gazda, H.T.; Eng, J.C.; Eichhorn, S.W.; Thiru, P.; Ghazvinian, R.; George, T.I.; Gotlib, J.R.; Beggs, A.H.; Sieff, C.A.; et al. Altered translation of GATA1 in Diamond-Blackfan anemia. Nature Medicine 2014, 20, 748–753. 10.1038/nm.3557.

60. Khajuria, R.K.; Munschauer, M.; Ulirsch, J.C.; Fiorini, C.; Ludwig, L.S.; McFarland, S.K.; Abdulhay, N.J.; Specht, H.; Keshishian, H.; Mani, D.R.; et al. Ribosome Levels Selectively Regulate Translation and Lineage Commitment in Human Hematopoiesis. Cell 2018, 173, 90–103.e19. 10.1016/j.cell.2018.02.036.

