## Supplementary material for "Novel 4D tensor decomposition-based approach integrating tri-omics profiling data can identify functionally relevant gene clusters": Supplementary_doc.pdf

Article

### Supplementary documement

**Y.-H. Taguchi [1,*] and Turki Turki [2]**

1 Department of Physics, Chuo University, Tokyo 112-8551, Japan
2 Department of Computer Science, King Abdulaziz University, Jeddah 21589, Saudi Arabia;

Received:
Revised:
Accepted:
Published:

**Copyright:** © 2026 by the authors. Submitted to *Journal Not Specified* for possible open access publication under the terms and conditions of the Creative Commons Attribution (CC BY) license.

#### 1. Understanding biological meaning

Usually, the biological meaning of the selected 1,781 genes are evaluated by enrichment analysis. Nevertheless, since enrichment within the selected genes is too much, we cannot

https://doi.org/10.3390/1010000

understand its meaning only using enrichment analysis. Thus, we make use of generative AI, although the following are confirmed by manual literature search, not based upon only the results of generative AI.

##### 1.1. Direct evaluation by generative AI

Direct evaluation by generative AI identified various functional gene subsets.

###### 1.1.1. Functional unit 1: Maintaining Genome Integrity and Controlling DNA Replication Licensing

DNA replication and the molecular machinery that maintains its stability. These play an essential role in redefining the dynamics of the cell cycle as cells transition from rapid proliferation to differentiation into specific tissues.

**Licensing of the MCM Complex and Replication Initiation** [1–3]: The list includes *Mcm*3, *Mcm*4, *Mcm*10 ans *Mcmbp*; the Minichromosome Maintenance (MCM) complex is a core apparatus that functions as a helicase in eukaryotic genome replication (Table 1).

| Gene symbol | Molecular function | Roles in biological process |
| --- | --- | --- |
| *Mcm*3 | Component of the MCM2-7 complex DNA helicase activity | Licensing of replication during the $G_1$ phase |
| *Mcm*4 | MCM2-7 Complex Core Subunit | DNA Double-Strand Dissociation and Replication Fork Progression |
| *Mcm*10 | Replication origin activator, MCM complex and Pol $\alpha$ binding helper | Initiation of replication and maintenance of genomic stability |
| *Mcmbp* | MCM Complex Chaperone and Disassembly Factor | Regulation at the Termination Stage of Replication and Cell Cycle Progression |

**Table 1.** Licensing of the MCM Complex and Replication Initiation [1–3]

**DNA Repair and Chromosome Segregation Fidelity**: Gene list includes *Rad*51, *Rad*54*b*, *Rad*54*l* which are homologous recombination modifier [4] and tumor suppressor genes such as BRCA2 [5]. These function to repair DNA damage that may occur during differentiation and maintain genomic integrity. Furthermore, it is rich in factors that govern the accurate distribution of chromosomes during mitosis. The presence of *Bub*1, *Bub*1*b*, *Cdc*20, *Cenpe*, *Cenpf*, *Cenph*, *Cenpk* indicates that the spindle assembly checkpoint (SAC) and kinetochore assembly are critical factors in the gene list. *Bub*1 and *Bub*1*b* detect abnormalities where chromosomes are not properly aligned and halt cell division at the spindle assembly checkpoint (SAC) [6]. *Cdc*20 serves as an essential activator of the Anaphase-Promoting Complex/Cyclosome (APC/C) [7]. However, until all chromosomes achieve proper bipolar attachment, the Spindle Assembly Checkpoint (SAC) prevents premature anaphase onset by sequestering into the Mitotic Checkpoint Complex (MCC) [8,9]. The *Cenph* and *Cenpk* genes identified in this analysis constitute the CCAN, the fundamental structure of the kinetochore [10], while *Cenpe* and *Cenpf* are responsible for monitoring the spindle assembly checkpoint (SAC) and the physical alignment of chromosomes [11]. The concentrated expression of these factors suggests that the gene list is highly organized as a functional unit ensuring the fidelity of cell division. *Bub*1*b*(*BubR*1), a member of the gene set, functions as a central component of the spindle assembly checkpoint (SAC). *Bub*1*b* monitors kinetochore dynamics until all chromosomes are correctly aligned at the spindle equatorial plane and appropriate tension is established [12]. When misaligned chromosomes are present, *Bub*1*b* forms the mitotic checkpoint complex (MCC) to inhibit the activity of the anaphase-promoting complex/cytokinase (APC/C), strictly suppressing

| Collagen Type | Primary Function | Related Diseases and Biological Significance |
|---|---|---|
| *Col1a1*/*Col1a2* | The primary structural component of bone, skin, and tendons | The most abundant fibrous collagen, providing tensile strength |
| *Col3a1* | Vascular wall, elastic maintenance of hollow organs | Causative gene for vascular-type Ehlers-Danlos syndrome (vEDS) |
| *Col4a1*/*Col4a2* | Formation of the basement membrane meshwork Cell scaffolding | Vascular stabilization |
| *Col5a1*/*Col5a2* | Regulation of Type I Collagen Fiber Diameter Control factors for fiber formation | Determining tissue quality |
| *Col6a1/2/3* | Microfibril formation | Cell adhesion A crucial link molecule bridging cells and the ECM |

**Table 2.** Collagen Networks [14]

the initiation of anaphase [8]. Therefore, *Bub1b* expression in this subset is thought to function as a 'guardian' ensuring the fidelity of chromosome distribution during cell fate transitions and maintaining genomic stability [13].

###### 1.1.2. Functional Unit 2: Renewal of the Extracellular Matrix (ECM) and the Basis of Tissue Morphogenesis

These are Matrisome-related factors that physically define the extracellular environment. This reflects the large-scale structural reorganization that occurs when cells form mesoderm-derived connective tissue and the vascular system.

**Collagen Networks and Matrix Diversity**: The list includes an extremely broad collagen family, such as *Col1a1*, *Col1a2*, *Col3a1*, *Col4a1*, *Col4a2*, *Col5a1*, *Col5a2*, *Col6a1*, *Col6a2*, *Col6a3*, *Col7a1*, *Col14a1*, *Col16a1*, and *Col23a1* (Table 2 [14]). In particular, *Col3a1* is essential for maintaining the integrity of blood vessels and fetal tissues, and its mutation can cause lethal arterial rupture. The simultaneous extraction of these diverse collagens in the gene list indicates that during specific differentiation stages—particularly the organization from LPM—it is not a single protein but the entire matrix "recipe" that is being renewed.

**Non-collagenous matrix components and modifying enzymes**: In addition to collagen, adhesion molecules such as *Postn* (periostin) [15], *Fbn1* (fibrillin-1) [16], *Lama2/4* (laminin), and *Nid1* (nidogen-1) [17], as well as *Lox* (lysyl oxidase), which cross-links the ECM to enhance its strength [18,19], have been identified. These factors function as "architects" that assemble secreted collagen into functional three-dimensional structures. Within the context of the lateral plate mesoderm, these units form the physical foundation of the future cardiovascular system, serous membranes, and connective tissue [20,21].

###### 1.1.3. Functional Unit 3: Mitochondrial Biogenesis and Energy Metabolism Conversion

As cells transition from undifferentiated stem cells to differentiated cells, the primary site of energy metabolism shifts from the cytoplasm (glycolysis) to mitochondria (oxidative phosphorylation). The gene list encompasses the molecular infrastructure required to execute this "metabolic reprogramming."

**Expansion of Mitochondrial Ribosomes (mitoribosomes)**: The list contains genes from the *Mrpl* (large subunit) and *Mrps* (small subunit) families at an astonishing density.

- *Mrpl11*, *Mrpl12*, *Mrpl15*, *Mrpl16*, *Mrpl24*, *Mrpl30*, *Mrpl33*, *Mrpl37*, *Mrpl45*, *Mrpl46*, *Mrpl47*, *Mrpl55*, and *Mrpl57*
- *Mrps11*, *Mrps16*, *Mrps23*, *Mrps26*, and *Mrps31*

These are ribosomal components specialized for protein synthesis within mitochondria and are essential for producing respiratory chain complex subunits encoded by the mitochondrial genome. Their accumulation in the gene list indicates that cells dramatically enhance mitochondrial "quality" and "quantity" during differentiation.

**Respiratory Chain Complex I and OXPHOS** : Furthermore, components and assembly factors of NADH:ubiquinone oxidoreductase (complex I), such as *Ndufa*7, *Ndufa*8, *Ndufa*9, *Ndufa*10, *Ndufaf*4, *Ndufaf*6, *Ndufb*10, *Ndufs*4, *Ndufs*8, and *Ndufv*2, are also major members of the gene list (Table 3). This unit symbolizes the "expansion project" of the mitochondrial power plant to meet the high energy demands required by differentiated cells.

| Gene | Functional Role | Clinical and Biological Significance |
|---|---|---|
| *Ndufa*10 | A component of Complex I, electron transport | Deficiencies cause mitochondrial encephalomyopathy and other conditions [22] |
| *Ndufaf*6 | Assembly cofactor for Complex I | Essential for normal respiratory chain assembly [23] |
| *Atp*5*k*/*Atp*5*j* | ATP synthase subunit | Responsible for ATP production, the final stage of OXPHOS [24] |

**Table 3.** Respiratory Chain Complex I and OXPHOS

###### 1.1.4. Functional Unit 4: Customization and Quality Control of the Protein Synthesis Apparatus

Optimizing the efficiency of intracellular protein production lines is essential to meet the demand for large quantities of new proteins (especially secreted proteins) that accompanies differentiation.

**Cytoplasmic Ribosomes and Control of Translation Initiation** : Numerous proteins from the *Rpl* and *Rps* systems (*Rpl*4, *Rpl*10*a*, *Rpl*21, *Rpl*22, *Rpl*24, *Rpl*32, *Rpl*34, *Rpl*35, *Rpl*37*a*, *Rps*10, *Rps*20, *Rps*24, *Rps*27, and *Rps*28) are listed. Ribosomal proteins are often thought to be stably expressed, but recent studies have highlighted "ribosomal heterogeneity," where changes in the composition of specific subunits selectively promote the translation of specific mRNA sets. Fluctuations in ribosomal factors in the gene list reflect the optimization of the translation machinery to generate a differentiation-specific proteome.

**Endoplasmic Reticulum (ER) Stress Response and Protein Secretion Pathways** : To support the aforementioned massive collagen secretion, the gene list also concentrates factors involved in protein quality control within the ER. These include chaperones such as *Hspa*5 (*BiP*), *Pdia*4, and *Uggt*1, as well as ER membrane transporters like *Tram*1, *Tram*2, *Sec*61*a*1, and *Sec*61*a*2 (Table 4) . This unit represents the logistical adjustments—expansion of ER capacity and enhanced quality control—that occur when cells shift into a "high-secretion mode."

###### 1.1.5. Functional Unit 5: Integration of Intracellular Transport and Signal Receptor Networks

Cell polarity formation and responsiveness to the external environment are dynamically regulated by the vesicular transport system.

**Rab GTPases and Vesicle Trafficking**: Rab family proteins such as *Rab*3*a*, *Rab*5*b*, *Rab*23, *Rab*25, *Rab*33*b*, and *Rab*43 are found in the gene list. In particular, *Rab*5*b* is the master regulator of early endosomes, governing the internalization of growth factor re-

| Gene | Function | Role in the Process |
|---|---|---|
| *Tram*1 | ER membrane translocation complex | Assists the translocation of polypeptides in translation into the ER lumen [25] |
| *Sec*61*a*1 | Core subunit of the translocon | Responsible for the ER entry of all secreted and membrane proteins [26] |
| *Hspa*5 | Major ER chaperone Promotes | Proper folding and senses ER stress [27] |

**Table 4.** Endoplasmic Reticulum (ER) Stress Response and Protein Secretion Pathways

ceptors and signal termination (or continuation) [28]. Furthermore, *Gga*2 and *Gga*3 are adaptors mediating cargo transport from the Golgi apparatus to endosomes, sorting specific cargo [29].

**Signal Transduction Pathways: TGF-$\beta$/BMP and Wnt Signaling**: The gene list also contains key signaling components that drive differentiation. TGF-$\beta$/BMP receptor families essential for mesoderm formation [8], such as *Tgfbr*1, *Tgfbr*3, *Tgfbrap*1, *Acvr*1, and *Acvr*2*a*, have been identified. These receptors govern the patterning of the lateral plate mesoderm and the final determination of cell fate. Furthermore, the presence of molecules such as *Fzd*1, *Fzd*7 (Wnt receptors) [30] and *Ctnna*1, *Ctnna*3 ($\alpha$-catenin) [31], which serve as interfaces between cell adhesion and signaling, suggests that physical interactions between cells modulate the sensitivity of signal reception.

###### 1.1.6. Functional Unit 6: Epigenetic Reprogramming and RNA Processing

The "interpretation" of gene expression within the nucleus is also highly regulated by the gene list.

**Histone Modification and Chromatin Structure Transformation**: Histone demethylases such as *Kdm*1*a*, *Kdm*1*b*, *Kdm*3*a*, *Kdm*4*a*, *Kdm*5*c* and *Ezh*1, *Ezh*2 (components of the Polycomb repressive complex 2) [32] are included. These function as "epigenetic switches" that repress (silence) the promoters of pluripotency-maintaining genes while making regions of differentiation-related genes accessible.

**Alternative splicing and RNA modifications**: The accumulation of splicing factors such as *Srsf*4, *Srsf*6, *Srsf*7, *Snrnp*27, *Snrpe*, qand *Snrpd*1 indicates that "proteome diversification" through splicing pattern switching [33], rather than merely changes in transcription levels, is a crucial aspect of the gene list during differentiation. Furthermore, the presence of RNA methyltransferases such as *Mettl*1, *Mettl*5, *Mettl*8, *Mettl*15, *Mettl*17 suggests that adjustments to mRNA stability and translation efficiency, mediated by modifications like m6A [34], are intensively occurring during critical junctures of cellular fate conversion.

###### 1.1.7. The gene list Depicts "Coupling of Morphogenesis and Energy Metabolism"

Beyond analyzing individual functional units (Fig. 1), examining how they integrate reveals that the gene list represents the physical implementation of biological "transitional states."

1. Synchronization of Structural Construction and Bioenergetics
   In the gene list, the simultaneous extraction of collagen (structure) and the mitochondrial apparatus (energy) is critically important. Collagen synthesis is an energy-intensive process requiring large amounts of oxygen, ATP, and $\alpha$-ketoglutarate (a metabolite of the TCA cycle). The gene list is thought to capture the synchrony of

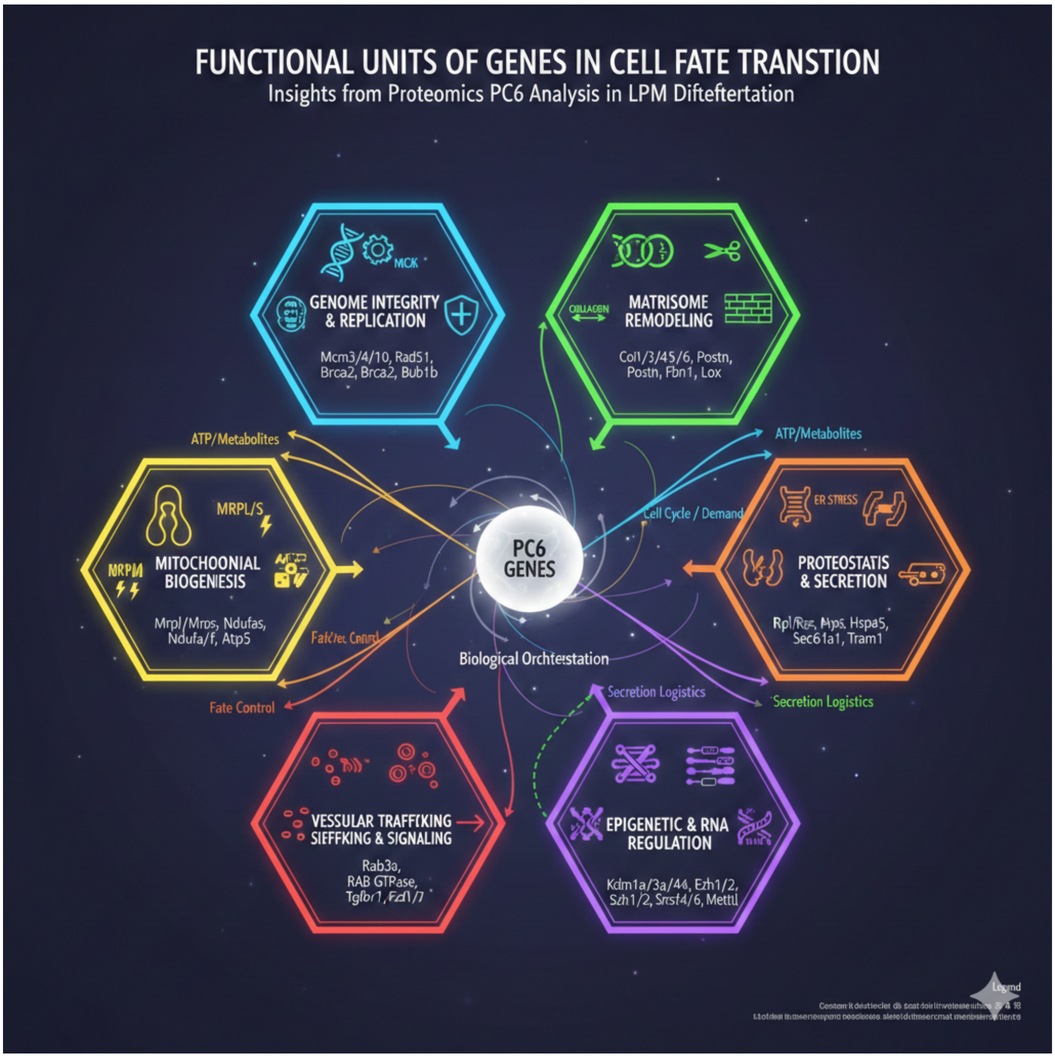

**Figure 1.** Functional units of genes in cell fate transition

a program based on economic rationality: cells simultaneously build the physical "walls of the tissue" while expanding the "power plants" to support it.

2. Redefining the Cell Cycle and Controlling the "Window" for Differentiation
   The inclusion of the *Mcm* gene cluster in the gene list may be closely related to the extension of the $G_1$ phase associated with differentiation. Embryonic stem cells possess an extremely short $G_1$ phase, but this phase lengthens during differentiation, allowing integration of differentiation signals. Since replication licensing by the MCM complex occurs during $G_1$, the behavior of *Mcm* genes in the gene list may represent a gear shift in the cell cycle engine to adjust the "window" for differentiation.
3. The gene list reveals the lateral plate mesoderm (LPM) signature.
   Comparing this information, these gene lists correspond to a specific phase of mouse LPM differentiation, particularly when progenitor cells lose epithelial polarity (involving *Ctnna*1 etc.), initiate migration, and deposit extensive extracellular matrix to form the scaffold for future connective tissue. The gene list is thus concluded to capture the proteomic fingerprint at this intersection of "Morphogenesis" and "Fate Commitment."

1.1.8. Conclusion

Analysis of the gene list revealed the existence of six major functional units during cell fate transition.

- **Genome Replication and Maintenance Unit**: *Mcm*3/4/10, homologous recombination, and mitotic checkpoint factors control the fidelity and dynamics of cell division during differentiation.
- **Matrix/Rearrangement Unit**: Constructs tissue-specific extracellular microenvironments via numerous collagens (*Col*1/3/4/5/6) and crosslinking enzymes (*Lox*).
- **Mitochondrial Biogenesis Unit**: Executes metabolic conversion from glycolysis to OXPHOS through expansion of mitochondrial ribosomes and respiratory chain complexes (NDUF).
- **Proteostasis/Secretion Unit**: Meets high secretory demands through ribosome customization, ER chaperones, and membrane translocation apparatus (Sec61).
- **Vesicle Transport/Signal Integration Unit**: Controls environmental responsiveness and cell polarity via Rab GTPases and TGF-$\beta$/Wnt receptors.
- **Epigenetic and RNA Regulation Unit**: Histone modification enzymes and splicing factors fundamentally rewrite the interpretation of gene expression.

These units do not function independently but achieve the complex task of maturing a single cell into a tissue component by complementing and coordinating with each other. The gene list scientifically extracts this biological "orchestration" in the form of distributed multi-layered omics.

##### *1.2. Analysis using generative AI after clustering enrichment analysis*

Generatie AI interpretation of the gene list through clustering enrichment analysis can depict another side of feature. The gene clusters analyzed formed multiple clusters exhibiting high enrichment scores, suggesting they constitute a highly interconnected network rather than merely independent pathways. In particular, functional units centered around ubiquitin-SUMO modification (Ubl modification) for protein quality control, genome stability maintenance linked to the cell cycle, bioenergetic metabolism originating in mitochondria and iron-sulfur cluster biosynthesis, and nutrient-responsive signaling (Hippo-TORC1) emerged as core pillars supporting cell survival and proliferation.

###### 1.2.1. Functional Unit 1: Integrated Control Unit for Genome Stability Maintenance and Cell Division

One of the most prominent functional units within this gene subset is the apparatus for maintaining genomic integrity and executing accurate cell division. This unit concentrates genes governing DNA replication, DNA damage response (DDR), chromosome segregation, and mitotic progression, revealing their extremely stringent temporal and spatial regulation.

**The Engine of DNA Replication: The CMG Helicase Complex**: The Cdc45-MCM2-7-GINS (CMG) complex plays a central role in the initiation and elongation of DNA replication [35]. This massive helicase complex, composed of 11 subunits, is responsible for dissociating double-stranded DNA at the replication fork during S phase. Formation of the CMG complex is tightly regulated by the cell cycle. After the MCM2-7 hexamer loads onto DNA during G1 phase, it becomes active in S phase through the recruitment of Cdc45 and the GINS tetramer (Sld5-Psf1-Psf2-Psf3).

Notably, cells preload 5 to 10 times more MCM hexamers than required to complete a normal S phase, known as "reserve MCMs." [36] These dormant origins function as a rescue mechanism when replication stress halts forks, enabling restart. This process is monitored by DNA damage response kinases such as ATR and ATM. When metabolic stress or DNA damage occurs, these reserve MCMs are activated, thereby preventing genomic instability before it arises.

**DNA Double-Strand Break (DSB) Repair Selection and the NuA4 Complex**: For double-strand breaks (DSBs), the most lethal form of DNA damage, cells employ two major repair pathways: non-homologous end joining (NHEJ) and homologous recombination (HR) [37]. This gene subset likely contains epigenetic modifiers that control the selection between these pathways (Table 5).

| Repair Pathway | Primary Cell Cycle | Characteristics | Key Involved Factors |
|---|---|---|---|
| Non-homologous end joining (NHEJ) | Overall cell cycle | Fast but prone to errors | 53BP1, Ku70/80 |
| Homologous recombination (HR) | S phase and G2 phase | High-precision repair using sister chromatids as templates | NuA4, BRCA1, Rad51 |

**Table 5.** DSB repair pathway

At the core of this process is the NuA4 (TIP60) histone acetyltransferase complex [38]. NuA4 is preferentially activated during S and G2 phases when sister chromatids are present, accumulating at damage sites via phosphorylation-dependent recruitment by the MRX (Mre11-Rad50-Xrs2) complex. By acetylating histones H4 and H2A, NuA4 maintains chromatin in an "open" state and physically inhibits the binding of 53BP1, which promotes NHEJ. This allows cells to select the error-prone HR pathway, enabling accurate repair of genomic information.

**Chromosome Segregation and Kinetochore Dynamics**: The success of cell division (mitosis) depends on whether replicated chromosomes are equally distributed to daughter cells. In this process, the kinetochore is a large protein complex formed at the centromere region of chromosomes, functioning as the mechanical interface connecting spindle microtubules to chromosomes.

Kinetochore complexes are broadly categorized into an inner layer (CCAN) present at the centromere throughout the cell cycle and an outer layer (KMN network) formed only during mitosis [36]. The Ndc80 complex in the outer layer directly binds to microtubules, generating traction to move chromosomes [39]. Additionally, Aurora B kinase, part of the chromosome passenger complex (CPC), corrects misassociations with microtubules and, in coordination with the spindle assembly checkpoint (SAC), delays the initiation of anaphase until all chromosomes are properly aligned [40].

###### 1.2.2. Functional Unit 2: Post-Translational Modification (PTM) Networks and Proteostasis Units

Within the gene clustering data, the highest enrichment score is observed for "Ubl conjugation" (Cluster 1, Score 23.53). This indicates that cells regulate protein states not merely by their presence or absence, but through "functional switching" via chemical modifications.

**The Ubiquitin-Proteasome System (UPS) and Specificity Determinants**: Ubiquitination is a process that adds ubiquitin molecules to target proteins through a three-step enzymatic reaction involving E1 (activation), E2 (conjugation), and E3 (ligation) enzymes. E3 ubiquitin ligases function as "specificity-determining factors" that recognize hundreds to thousands of target proteins, forming the cornerstone of cellular protein metabolism (proteostasis) [41].

This dataset contains numerous proteins that include WD40 repeats and Kelch repeats, which play a crucial role as substrate recognition adaptors in the Cullin-RING ligase (CRL) complex.

- **WD40 repeat**: Forms a $\beta$-propeller structure, serving as a robust platform for protein-protein interactions. For example, the CRL4-DCAF1 complex recruits DNA repair factors and cell cycle regulators (such as Cdt1) via WD40 proteins, inducing their degradation [42].

- **Kelch repeat**: Similarly possesses a $\beta$-propeller structure and functions as an adapter for Cullin 3 (Cul3)-type ligases. Specifically, KLHL21 targets Aurora B during mitosis, ensuring proper progression of cytokinesis [43].

**SUMOylation and STUbL-Mediated Complex Regulatory Circuits** : SUMOylation (Small Ubiquitin-like Modifier) is a modification that regulates the localization and interactions of target proteins. When combined with ubiquitination, it forms the more sophisticated "SUMO Targeted Ubiquitin Ligase (STUbL)" pathway [44].

STUbLs such as RNF4 and RNF111 recognize poly-SUMOylated proteins via their SUMO interaction motif (SIM) and add ubiquitin chains to them, leading to degradation by the proteasome [45]. This mechanism is critically important in the stress response during DNA replication. When the replication fork halts, many replicasome components become SUMOylated. This signals STUbL to act, "cleaning up" old proteins and creating space for repair enzymes to access the DNA. Thus, the synergy between SUMOylation and ubiquitination functions as a logic circuit enabling cells to respond rapidly and irreversibly to critical situations.

**Maintenance of Reversibility by Deubiquitinating Enzymes (DUBs)**: The dynamics of PTM units are determined not only by ligase-mediated modification but also by the "erasure" process carried out by deubiquitinating enzymes (DUBs) and SUMO-specific proteases (SENPs) [46]. DUBs, such as the USP (Ubiquitin-Specific Protease) family, play a role in preventing premature protein degradation and maintaining the cellular pool of free ubiquitin by removing unwanted ubiquitin chains. This constant balance (turnover) between modification and demodification enables the instantaneous switching of cell cycle progression and the "on-off" control of signal transduction.

###### 1.2.3. Functional Unit 3:Mitochondrial Bioenergetics and Iron-Sulfur Cluster Biosynthesis Unit

One intriguing finding in this analysis is that mitochondrial-associated genes form functional units that closely interact with nuclear DNA metabolic factors.

**Construction of Respiratory Chain Complex I and Respiratory Supercomplex**: Respiratory chain complex I (NADH:ubiquinone oxidoreductase), located in the mitochondrial inner membrane, is a massive enzyme complex responsible for initiating the electron transport chain, composed of over 45 subunits [47]. This complex pumps protons from the matrix to the intermembrane space during the transfer of electrons from NADH to coenzyme Q10, thereby establishing the electrochemical potential that drives ATP synthesis.

The assembly of Complex I requires more than 24 assembly factors, and this process is closely synchronized with the insertion of iron-sulfur (Fe-S) clusters [48]. Recent studies have revealed that the ISD complex, containing cystathionine synthase (Nfs1), the central enzyme in Fe-S cluster synthesis, directly interacts with respiratory chain supercomplexes. This suggests that cells spatially regulate the supply of cofactors required for electron transport to match their energy production capacity (respiratory activity).

**Iron-Sulfur Cluster Biosynthesis: A Lifeline for Nuclear Function**: The true importance of mitochondria extends far beyond being mere "powerhouses." The mitochondrial iron-sulfur cluster (ISC) assembly apparatus is the sole source for the maturation of Fe-S proteins throughout the cell [49]. Intermediates synthesized within mitochondria (X-S compounds) are exported to the cytoplasm via the ABC transporter ABCB7 (Atm1 in yeast) and handed over to the cytoplasmic iron-sulfur protein assembly (CIA) machinery for incorporation into final target proteins [50].

Remarkably, many enzymes governing genomic stability within the nucleus require clusters as the foundation of their activity.

- **DNA replication**: DNA primase and DNA polymerases $\alpha$, $\delta$, and $\epsilon$ all possess Fe-S clusters [51]. In DNA primase, the oxidation state of the Fe-S cluster functions as a "switch" for DNA binding, controlling the initiation of primer synthesis [52].
- **DNA Repair and Helicases**: Helicase enzymes such as DNA2, FANCJ, and XPD that dissociate DNA also depend on Fe-S clusterss [51]. Deficiency of these clusters leads to replication fork stalling and accumulation of DNA damage, causing severe genomic instability disorders such as Fanconi anemia and xeroderma pigmentosum.

Thus, mitochondrial metabolic dysfunction goes beyond mere energy deficiency to directly disrupt the nuclear genome maintenance function. This mitochondrial-nuclear coordination unit can be considered one of the fundamental principles of life, directly linking the cell's "energy state" to the "preservation of genetic information."

###### 1.2.4. Functional Unit 4: Proliferation Control and Transcription Integration Signaling Unit

The decision of when a cell grows and when it divides is made by integrating signals from the environment, such as nutrients and physical contact stimuli, with intrinsic clock mechanisms. This subset of genes includes the Hippo pathway, mTORC1 pathway, and mediator complex, which are responsible for this integrated decision-making.

**The Hippo Pathway and mTORC1 Crossroads**: The Hippo signaling pathway plays a central role in controlling organ size and maintaining tissue homeostasis, primarily by regulating cell number (proliferation and apoptosis) [53]. Meanwhile, mTORC1 (mechanistic target of rapamycin complex 1) controls cell size (protein synthesis and ribosome biogenesis) in response to nutritional and energy states [54].

The key finding in this report is the direct phosphorylation-dependent crosstalk between these two major growth control pathways. LATS1/2, the core kinase of the Hippo pathway, directly phosphorylates Raptor at Ser606, an essential component of mTORC1 [53]. This phosphorylation modification inhibits the interaction between Raptor and its activator, the Rheb GTPase, thereby attenuating mTORC1 activity.

- **During Hippo pathway activation (e.g., high cell density)**: LATS1/2 activates $\rightarrow$ Raptor Ser606 is phosphorylated $\rightarrow$ mTORC1 is inhibited $\rightarrow$ Cell growth stops [55].
- **When Hippo pathway is inactivated (e.g., tissue injury)**: LATS1/2 becomes inactive $\rightarrow$ Raptor remains unphosphorylated $\rightarrow$ mTORC1 activates $\rightarrow$ Rapid cell hypertrophy and proliferation commence [53].

Through this mechanism, cells simultaneously calculate "spatial constraints (Hippo" and "resource availability (mTORC1)" to derive an appropriate growth strategy.

**Circadian Rhythm-Mediated Gate Control of the Cell Cycle**: Cellular processes are also closely synchronized with the 24-hour circadian rhythm [56]. The central clock (CLOCK/BMAL1) rhythmically regulates the expression of key genes controlling the cell cycle [57]. For example, the promoter of WEE1, a kinase that inhibits mitosis, contains an E-box sequence to which BMAL1 binds. This mechanism "gates" (restricts) the timing of entry into mitosis to specific periods of the day [58].

Conversely, the state of the cell cycle also provides feedback to the clock mechanism. CDK1, the master regulator of cell division, phosphorylates REV-ERB$\alpha$, an inhibitor of the circadian clock, thereby promoting its degradation by the E3 ligase FBXW7 [58]. This process resets the clock in conjunction with cell division. This reciprocal control unit enables cells to adopt an evolutionarily optimal survival strategy: performing DNA replication during periods of high energy metabolism and completing division during periods of low environmental risk, such as reduced ultraviolet radiation [56].

###### 1.2.5. Functional Unit 5: Dynamic Coordination Between the Metabolic-Epigenomic Axis and Nuclear Functions

Among the gene subsets are groups involved in one-carbon metabolism and methylation reactions, forming a “metabolic-epigenomic integration unit” that translates the cell’s nutritional state into the physical structure of the genome [59].

**One-Carbon Metabolism and SAM Supply: The Source of Methylation**: One-carbon metabolism, comprising the folate cycle and methionine cycle, generates S-adenosylmethionine (SAM), the sole methyl donor within cells [59]. SAM serves as the substrate for DNA methyltransferases (DNMTs) and histone methyltransferases (HMTs), constituting the absolute limiting factor for epigenetic “writing.” [60]

Dietary intake of vitamin B12 and methionine fluctuates blood SAM levels, which directly influences histone methylation states (activation marks like H3K4me3 and suppression marks like H3K9/K27me3) [61]. During nutritional deficiency, the flux of one-carbon metabolism is preferentially allocated to maintaining SAM levels, activating an adaptive mechanism that strives to preserve heterochromatin and safeguard the expression patterns of essential genes [60].

**Metabolic Reprogramming in the DNA Damage Response**: When DNA damage occurs, cells not only repair it but also dramatically alter central metabolism to support the repair process [62]. The damage-sensing checkpoint kinase (Mec1/Tel1) SUMO-modifies Snf1 (mammalian AMPK), a master regulator of energy metabolism, thereby suppressing its activity [63].

Inhibition of Snf1 reduces mitochondrial respiration and shifts energy metabolism toward glycolysis and fermentation (the so-called Warburg effect) [63]. This metabolic shift is thought to be a "defensive reprogramming" to minimize the generation of reactive oxygen species (ROS), which could cause further DNA damage during the repair process [63]. Thus, the metabolic control unit serves not merely as an energy source, but as a "foundation" for successfully executing the costly and delicate process of genome repair.

Table 6 summarizes the correspondence between above five functional units and 77 gene clusters.

###### 1.2.6. Cellular Architecture and Organelle Dynamics Unit

The final pillar supporting the functional units of the cell is the module responsible for physical structural maintenance and the precise inheritance of organelles [62].

**FERM Domain Proteins: Interface Between Membrane and Cytoskeleton**: The FERM (Four-point-one, Ezrin, Radixin, Moesin) domain serves as a universal platform anchoring the cytoskeleton to the cell membrane [64]. Proteins containing the FERM domain, particularly Merlin (NF2) and Expanded (Ex), function as "membrane sensors" at the very top of the Hippo pathway [65].

Merlin recruits LATS1/2 kinases to the cell membrane via adaptors such as the RASSF family when it senses cell-cell contacts or membrane tension. This proximity effect near the membrane ignites the kinase cascade, ultimately blocking the nuclear translocation of the growth factors YAP/TAZ [66]. This functional unit acts as a translator, converting the cell’s surrounding "physical environment" (whether neighboring cells are present or the substrate is rigid) into a molecular signal for proliferation arrest [67].

**The Golgi apparatus’s “mitotic checkpoint” and inheritance**: During cell division, large organelles such as the endoplasmic reticulum (ER) and Golgi apparatus undergo drastic morphological changes to ensure equal distribution to daughter cells [68]. The Golgi apparatus, in particular, disassembles its ribbon-like structure during the G2/M transition phase, forming independent stacks that further fragment into minute vesicles and diffuse throughout the cytoplasm [69].

| | |
|---|---|
| Functional Unit 1: Integrated Control Unit for Genome Stability Maintenance and Cell Division | |
| This core group integrates and controls the entire cycle from DNA replication and damage repair to the actual physical cell division (chromosome segregation). | |
| Cluster 4, 14, 27, 31, 44, 54 | DNA repair, DNA damage response, homologous recombination, NuA4 complex, topoisomerase |
| Cluster 11, 57 | DNA Replication · CMG Complex (Cdc45-MCM-GINS) |
| Cluster 2, 51, 52 | Progression of Cell Division and Mitosis |
| Cluster 5, 22 | Kinetochore Chromatin Passenger Complex (CPC) Spindle pole proteins |
| Cluster 9, 12 | Microtubule-binding kinesin motor |
| Functional Unit 2: Post-Translational Modification (PTM) Networks and Proteostasis Units | |
| This network regulates protein lifespan and function through the dynamics of "addition (ligase)" and "removal (DUB/SENP)" in ubiquitination and SUMOylation. | |
| Cluster 1, 35 | SUMO2 Modification · SUMOylation (Mms21, etc.) |
| Cluster 7, 15, 69 | Protein ubiquitination, RING-type ligase, protein degradation |
| Cluster 28, 45 | WD40 Repeat · Kelch Repeat (E3 ligase substrate recognition adaptor) |
| Cluster 66 | Deubiquitination (elimination process: DUBs) |
| Cluster 77 | De-SUMOylation (Elimination Process: SENPs) |
| Functional Unit 3:Mitochondrial Bioenergetics and Iron-Sulfur Cluster Biosynthesis Unit | |
| It is a specialized unit centered on mitochondria that supplies essential cofactors (Fe-S) for cellular energy production (respiration) and for DNA replication and repair enzymes within the nucleus. | |
| Cluster 3, 56 | Mitochondrial outer membrane, matrix, and electron transport chain (respiratory chain) |
| Cluster 17, 30, 40 | Iron-Sulfur (Fe-S) Cluster Bonding and Assembly |
| Functional Unit 4: Proliferation Control and Transcription Integration Signaling Unit | |
| This signaling pathway detects physical environments such as cell density (Hippo) and nutritional status (mTORC1), integrating this information into final gene expression (transcription). | |
| Cluster 26 | FERM domain (connects the cell membrane to the cytoskeleton and senses contact inhibition) |
| Cluster 34 | Hippo Signaling Pathway (Translator of Cell Density and Contact Inhibition) |
| Cluster 36 | mTORC1/TORC1 signaling (nutrient and amino acid sensing) |
| Cluster 32 | Mediator Complex (Integrating Upstream Signals into Transcription) |
| Cluster 74 | Mitochondrial outer membrane and matrix |
| Functional Unit 5: Dynamic Coordination Between the Metabolic-Epigenomic Axis and Nuclear Functions | |
| These are a group of signals that detect the physical environment (cell density) and nutritional status, thereby determining whether to turn proliferation on or off. | |
| Cluster 50 | One-carbon metabolism and folate pool (supply of DNA synthesis materials and S-adenosylmethionine) |
| Cluster 41 | Methyl Transferase · Histone Methylation (Epigenome Regulation) |
| Cluster 6, 19 | Endoplasmic Reticulum (ER) and Golgi Apparatus Dynamics |
| Cluster 10, 18 | Intracellular transport of proteins and lysosomes (mTORC1 scaffold) |
| Cluster 13 | Ribosome and Translation System |

**Table 6.** Correspondence between five functional unit and 77 gene clusters.

It has been revealed that this disassembly process itself functions as the "Golgi checkpoint" that permits the progression of mitosis [68]. If Golgi disassembly is incomplete, the activation of mitotic kinases such as Aurora A is delayed, inhibiting spindle formation [69]. Furthermore, Golgi proteins such as GM130 and the GRASP family actively participate in organizing spindle microtubules during mitosis, shifting away from their role in membrane transport [70]. This demonstrates that the cell temporarily changes the Golgi apparatus's function from a "transportation unit" to a "structural maintenance unit" during mitosis, symbolizing the flexibility and rationality of functional units.

###### 1.2.7. Integrated Analysis of Higher-Order Crosstalk and Causality Among Functional Units

The individual functional units described thus far do not exist independently. Instead, they form a vast integrated system for sustaining life through specific "hub" proteins and metabolites.

**The Domino Effect of Genome Maintenance Originating from the Mitochondrial ISC Apparatus**: One typical causal chain extends from the mitochondrial bioenergetic unit to the nuclear genome stability unit. When mitochondrial iron-sulfur cluster synthesis (ISC) apparatus becomes dysfunctional due to nutritional deficiencies or genetic factors, cofactor supply to nuclear polymerase $\epsilon$ and DNA helicase II ceases [52]. This destabilizes replication fork progression (replication stress) and triggers excessive consumption of dormant replication origins (pre-MCM complexes) [71].

When replication stress exceeds a threshold, ATM/ATR kinase is activated, leading to the recruitment of NuA4 acetyltransferase to the damaged site [72]. Simultaneously, metabolic units suppress respiration via Snf1/AMPK SUMOylation to ensure safety during repair work1 [63]. Thus, mitochondria—functional units seemingly distant from the nucleus—govern the most fundamental information replication process within the nucleus.

**Signal "Switching" and Temporal Control via PTM Modification**: Another unifying trend is that PTM (Ubl modification) units determine the "temporal switching (timing)" of all other functional units. The irreversibility of the cell cycle is ensured not only by reversible modifications like phosphorylation but also by "one-way disposal" via ubiquitin-mediated proteasome degradation [73].

The integration of Hippo signaling and mTORC1 also leads to sustained phenotypic changes not only through phosphorylation but also by promoting the degradation of clock proteins and growth factors via F-box proteins (such as FBXW7) [57]. The degradation of SUMOylated proteins by STUbL also functions as a "cleanup" after DNA repair is complete, enabling the transition to the next step by rapidly removing tools once they have served their purpose [74].

###### 1.2.8. Conclusion: The Significance of Gene Subsets as Life Systems

The conclusion drawn from the analysis of the provided gene subsets is that a cell is a "Dynamic equilibrium system that is both closed and open, wherein each functional unit utilizes the outputs of others as inputs." (Fig. 2)

1. The genome stability unit ensures information integrity, but its operation requires Fe-S clusters and ATP supplied by the mitochondrial metabolism unit.
2. The PTM/proteostasis unit determines the system's execution speed and directionality, translating signals from the proliferation control unit into specific changes in protein lifespan.
3. The Epigenome-Metabolism Axis reflects the extracellular nutrient environment into the "settings" for medium-to-long-term gene expression, fine-tuning the overall system output (proliferation or quiescence).

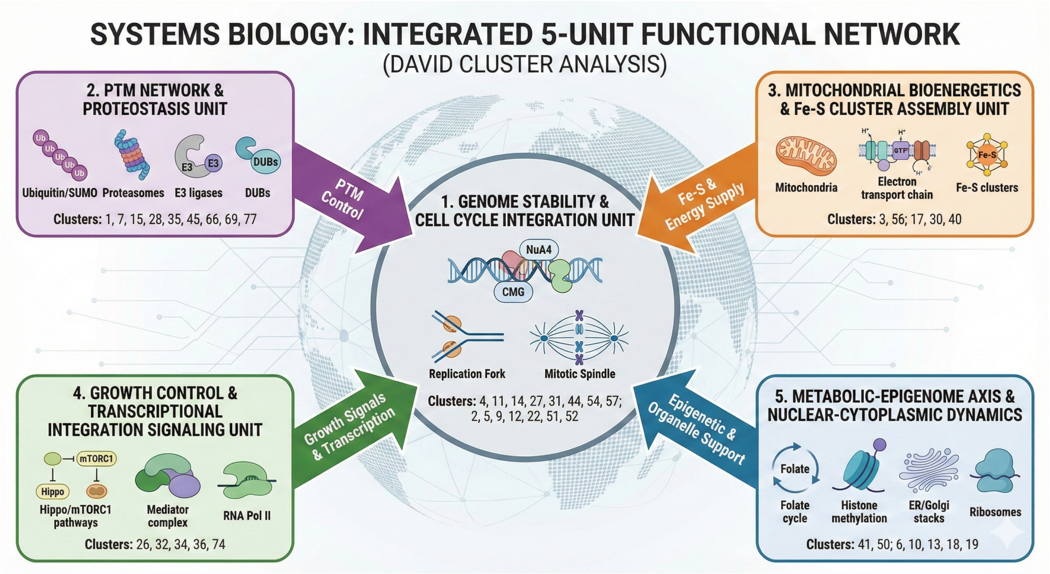

**Figure 2.** Five functional units in this section.

4. The Cell Architecture Unit provides the physical "space" where all these biochemical reactions occur, translating spatial information into biochemical language.

#### References

1. Matson, J.P.; Dumitru, R.; Coryell, P.; Baxley, R.M.; Chen, W.; Twaroski, K.; Webber, B.R.; Tolar, J.; Bielinsky, A.K.; Purvis, J.E.; et al. Rapid DNA replication origin licensing protects stem cell pluripotency. *eLife* **2017**, *6*, e30473. https://doi.org/10.7554/eLife.30473.
2. Douglas, M.E.; Diffley, J.F. Recruitment of Mcm10 to Sites of Replication Initiation Requires Direct Binding to the Minichromosome Maintenance (MCM) Complex*. *Journal of Biological Chemistry* **2016**, *291*, 5879–5888. https://doi.org/https://doi.org/10.1074/jbc.M115.707802.
3. Nishiyama, A.; Frappier, L.; Méchali, M. MCM-BP regulates unloading of the MCM2–7 helicase in late S phase. *Genes & Development* **2011**, *25*, 165–175, [https://genesdev.cshlp.org/content/25/2/165.full.pdf+html]. https://doi.org/10.1101/gad.614411.
4. Tavares, E.M.; Wright, W.D.; Heyer, W.D.; Cam, E.L.; Dupaigne, P. In vitro role of Rad54 in Rad51-ssDNA filament-dependent homology search and synaptic complexes formation. *Nature Communications* **2019**, *10*, 4058. https://doi.org/10.1038/s41467-019-12082-z.
5. Gorodetska, I.; Kozeretska, I.; Dubrovska, A. <i>BRCA</i> Genes: The Role in Genome Stability, Cancer Stemness and Therapy Resistance. *J Cancer* **2019**, *10*, 2109–2127. https://doi.org/10.7150/jca.30410.
6. Bolanos-Garcia, V.M.; Blundell, T.L. BUB1 and BUBR1: multifaceted kinases of the cell cycle. *Trends in Biochemical Sciences* **2011**, *36*, 141–150. https://doi.org/https://doi.org/10.1016/j.tibs.2010.08.004.
7. Visintin, R.; Prinz, S.; Amon, A. *CDC20* and *CDH1*: A Family of Substrate-Specific Activators of APC-Dependent Proteolysis. *Science* **1997**, *278*, 460–463, [https://www.science.org/doi/pdf/10.1126/science.278.5337.460]. https://doi.org/10.1126/science.278.5337.460.
8. Sudakin, V.; Chan, G.K.; Yen, T.J. Checkpoint inhibition of the APC/C in HeLa cells is mediated by a complex of BUBR1, BUB3, CDC20, and MAD2. *Journal of Cell Biology* **2001**, *154*, 925–936, [https://rupress.org/jcb/article-pdf/154/5/925/1512753/jcb1545925.pdf]. https://doi.org/10.1083/jcb.200102093.
9. Hwang, L.H.; Lau, L.F.; Smith, D.L.; Mistrot, C.A.; Hardwick, K.G.; Hwang, E.S.; Amon, A.; Murray, A.W. Budding Yeast Cdc20: A Target of the Spindle Checkpoint. *Science* **1998**, *279*, 1041–1044, [https://www.science.org/doi/pdf/10.1126/science.279.5353.1041]. https://doi.org/10.1126/science.279.5353.1041.

10. Okada, M.; Cheeseman, I.M.; Hori, T.; Okawa, K.; McLeod, I.X.; Yates, J.R.; Desai, A.; Fukagawa, T. The CENP-H-I complex is required for the efficient incorporation of newly synthesized CENP-A into centromeres. *Nature cell biology* **2006**, *8*, 446—457. https://doi.org/10.1038/ncb1396.
11. Yao, X.; Abrieu, A.; Zheng, Y.; Sullivan, K.F.; Cleveland, D.W. CENP-E forms a link between attachment of spindle microtubules to kinetochores and the mitotic checkpoint. *Nature Cell Biology* **2000**, *2*, 484–491. https://doi.org/10.1038/35019518.
12. Lampson, M.A.; Kapoor, T.M. The human mitotic checkpoint protein BubR1 regulates chromosome–spindle attachments. *Nature Cell Biology* **2005**, *7*, 93–98. https://doi.org/10.1038/ncb1208.
13. Baker, D.J.; Jeganathan, K.B.; Cameron, J.D.; Thompson, M.; Juneja, S.; Kopecka, A.; Kumar, R.; Jenkins, R.B.; de Groen, P.C.; Roche, P.; et al. BubR1 insufficiency causes early onset of aging-associated phenotypes and infertility in mice. *Nature Genetics* **2004**, *36*, 744–749. https://doi.org/10.1038/ng1382.
14. Ricard-Blum, S. The Collagen Family. *Cold Spring Harbor Perspectives in Biology* **2011**, *3*, [http://cshperspectives.cshlp.org/content/3/1/a004978.full.pdf+html]. https://doi.org/10.1101/cshperspect.a004978.
15. Conway, S.J.; Izuhara, K.; Kudo, Y.; Litvin, J.; Markwald, R.; Ouyang, G.; Arron, J.R.; Holweg, C.T.J.; Kudo, A. The role of periostin in tissue remodeling across health and disease. *Cellular and Molecular Life Sciences* **2014**, *71*, 1279–1288. https://doi.org/10.1007/s00018-013-1494-y.
16. Ramirez, F.; Sakai, L.Y. Biogenesis and function of fibrillin assemblies. *Cell and Tissue Research* **2009**, *339*, 71. https://doi.org/10.1007/s00441-009-0822-x.
17. Yurchenco, P.D. Basement Membranes: Cell Scaffoldings and Signaling Platforms. *Cold Spring Harbor Perspectives in Biology* **2011**, *3*, [http://cshperspectives.cshlp.org/content/3/2/a004911.full.pdf+htm https://doi.org/10.1101/cshperspect.a004911.
18. Lucero, H.A.; Kagan, H.M. Lysyl oxidase: an oxidative enzyme and effector of cell function. *Cellular and Molecular Life Sciences* **2006**, *63*, 2304–2316. https://doi.org/10.1007/s00018-006-6149-9.
19. Mäki, J.M.; Räsänen, J.; Tikkanen, H.; Sormunen, R.; Mäkikallio, K.; Kivirikko, K.I.; Soininen, R. Inactivation of the Lysyl Oxidase Gene <i>Lox</i> Leads to Aortic Aneurysms, Cardiovascular Dysfunction, and Perinatal Death in Mice. *Circulation* **2002**, *106*, 2503–2509, [https://www.ahajournals.org/doi/pdf/10.1161/01.CIR.0000038109.84500.1E]. https://doi.org/10.1161/01.CIR.0000038109.84500.1E.
20. Rozario, T.; DeSimone, D.W. The extracellular matrix in development and morphogenesis: A dynamic view. *Developmental Biology* **2010**, *341*, 126–140. Special Section: Morphogenesis, https://doi.org/https://doi.org/10.1016/j.ydbio.2009.10.026.
21. Gilbert, S.F.; Barresi, M.J.F. *Developmental Biology*, 11th ed.; Sinauer Associates, 2016.
22. Hoefs, S.J.G.; Rodenburg, R.J.; Smeitink, J.A.M.; van den Heuvel, L.P. NDUFA10 mutations cause complex I deficiency in a patient with Leigh disease. *European Journal of Human Genetics* **2011**, *19*, 270–274. https://doi.org/10.1038/ejhg.2010.204.
23. Valentino, M.L.; et al. NDUFAF6 mutations are a new cause of Leigh syndrome. *American Journal of Human Genetics* **2012**, *91*, 154–161.
24. Jonckheere, A.I.; Smeitink, J.A.M.; Rodenburg, R.J.T. Mitochondrial ATP synthase: architecture, function and pathology. *Journal of Inherited Metabolic Disease* **2012**, *35*, 211–225. https://doi.org/10.1007/s10545-011-9382-9.
25. Görlich, D.; Rapoport, T.A. Protein translocation into proteoliposomes reconstituted from purified components of the endoplasmic reticulum membrane. *Cell* **1993**, *75*, 615–630. https://doi.org/https://doi.org/10.1016/0092-8674(93)90483-7.
26. Robeson, L.; Casanova-Morales, N.; Burgos-Bravo, F.; Alfaro-Valdés, H.M.; Lesch, R.; Ramírez-Álvarez, C.; Valdivia-Delgado, M.; Vega, M.; Matute, R.A.; Schekman, R.; et al. Characterization of the interaction between the Sec61 translocon complex and ppαF using optical tweezers. *Protein Science* **2024**, *33*, e4996, [https://onlinelibrary.wiley.com/doi/pdf/10.1002/pro.4996]. https://doi.org/https://doi.org/10.1002/pro.4996.
27. Wang, J.; Lee, J.; Liem, D.; Ping, P. HSPA5 Gene encoding Hsp70 chaperone BiP in the endoplasmic reticulum. *Gene* **2017**, *618*, 14–23. https://doi.org/https://doi.org/10.1016/j.gene.2017.03.005.

28. Zerial, M.; McBride, H. Rab proteins as membrane organizers. *Nature Reviews Molecular Cell Biology* **2001**, *2*, 107–117. https://doi.org/10.1038/35052055.
29. Boman, A.L. GGA proteins: new players in the sorting game. *Journal of Cell Science* **2001**, *114*, 3413–3418, [https://journals.biologists.com/jcs/article-pdf/114/19/3413/1357062/3413.pdf]. https://doi.org/10.1242/jcs.114.19.3413.
30. Kim, M.; Lee, H.C.; Tsedensodnom, O.; Hartley, R.; Lim, Y.S.; Yu, E.; Merle, P.; Wands, J.R. Functional interaction between Wnt3 and Frizzled-7 leads to activation of the Wnt/$\beta$-catenin signaling pathway in hepatocellular carcinoma cells. *Journal of Hepatology* **2008**, *48*, 780–791. https://doi.org/https://doi.org/10.1016/j.jhep.2007.12.020.
31. Sehgal, R.N.; Gumbiner, B.M.; Reichardt, L.F. Antagonism of Cell Adhesion by an $\alpha$-Catenin Mutant, and of the Wnt-signaling Pathway by $\alpha$-Catenin in Xenopus Embryos. *Journal of Cell Biology* **1997**, *139*, 1033–1046, [https://rupress.org/jcb/article-pdf/139/4/1033/1487530/29227.pdf]. https://doi.org/10.1083/jcb.139.4.1033.
32. Gray, Z.H.; Honer, M.A.; Ghatalia, P.; Shi, Y.; Whetstine, J.R. 20 years of histone lysine demethylases: From discovery to the clinic and beyond. *Cell* **2025**, *188*, 1747–1783. https://doi.org/https://doi.org/10.1016/j.cell.2025.02.023.
33. Li, D.; Yu, W.; Lai, M. Towards understandings of serine/arginine-rich splicing factors. *Acta Pharmaceutica Sinica B* **2023**, *13*, 3181–3207. https://doi.org/https://doi.org/10.1016/j.apsb.2023.05.022.
34. Tooley, J.G.; Catlin, J.P.; Tooley, C.E.S. METTLing in Stem Cell and Cancer Biology. *Stem Cell Reviews and Reports* **2023**, *19*, 76–91. https://doi.org/10.1007/s12015-022-10444-7.
35. Georgescu, R.; Yuan, Z.; Bai, L.; de Luna Almeida Santos, R.; Sun, J.; Zhang, D.; Yurieva, O.; Li, H.; O'Donnell, M.E. Structure of eukaryotic CMG helicase at a replication fork and implications to replisome architecture and origin initiation. *Proceedings of the National Academy of Sciences* **2017**, *114*, E697–E706, [https://www.pnas.org/doi/pdf/10.1073/pnas.1620500114]. https://doi.org/10.1073/pnas.1620500114.
36. Xiang, S.; Reed, D.R.; Alexandrow, M.G. The CMG helicase and cancer: a tumor "engine" and weakness with missing mutations. *Oncogene* **2023**, *42*, 473–490. https://doi.org/10.1038/s41388-022-02572-8.
37. Clouaire, T.; Legube, G. DNA double strand break repair pathway choice: a chromatin based decision? *Nucleus* **2015**, *6*, 107–113, [https://doi.org/10.1080/19491034.2015.1010946]. PMID: 25675367, https://doi.org/10.1080/19491034.2015.1010946.
38. Pollina, E.A.; Gilliam, D.T.; Landau, A.T.; Lin, C.; Pajarillo, N.; Davis, C.P.; Harmin, D.A.; Yap, E.L.; Vogel, I.R.; Griffith, E.C.; et al. A NPAS4–NuA4 complex couples synaptic activity to DNA repair. *Nature* **2023**, *614*, 732–741. https://doi.org/10.1038/s41586-023-05711-7.
39. Powers, A.F.; Franck, A.D.; Gestaut, D.R.; Cooper, J.; Gracyzk, B.; Wei, R.R.; Wordeman, L.; Davis, T.N.; Asbury, C.L. The Ndc80 Kinetochore Complex Forms Load-Bearing Attachments to Dynamic Microtubule Tips via Biased Diffusion. *Cell* **2009**, *136*, 865–875. https://doi.org/https://doi.org/10.1016/j.cell.2008.12.045.
40. Carmena, M.; Wheelock, M.; Funabiki, H.; Earnshaw, W.C. The chromosomal passenger complex (CPC): from easy rider to the godfather of mitosis. *Nature Reviews Molecular Cell Biology* **2012**, *13*, 789–803. https://doi.org/10.1038/nrm3474.
41. Fletcher, A.J.; Mabbitt, P.D. Editorial: Reviews in ubiquitin signaling: 2022. *Frontiers in Molecular Biosciences* **2023**, *Volume 10 - 2023*. https://doi.org/10.3389/fmolb.2023.1275393.
42. Xu, C.; Min, J. Structure and function of WD40 domain proteins. *Protein & Cell* **2011**, *2*, 202–214, [https://academic.oup.com/proteincell/article-pdf/2/3/202/48344557/13238_2011_article_1018.pdf]. https://doi.org/10.1007/s13238-011-1018-1.
43. Adams, J.; Kelso, R.; Cooley, L. The kelch repeat superfamily of proteins: propellers of cell function. *Trends in Cell Biology* **2000**, *10*, 17–24. https://doi.org/10.1016/S0962-8924(99)01673-6.
44. Zhou, H.; Deng, N.; Li, Y.; Hu, X.; Yu, X.; Jia, S.; Zheng, C.; Gao, S.; Wu, H.; Li, K. Distinctive tumorigenic significance and innovative oncology targets of SUMOylation. *Theranostics* **2024**, *14*, 3127–3149. https://doi.org/10.7150/thno.97162.
45. Han, J.; Mu, Y.; Huang, J. Preserving genome integrity: The vital role of SUMO-targeted ubiquitin ligases. *Cell Insight* **2023**, *2*, 100128. https://doi.org/https://doi.org/10.1016/j.cellin.2023.100128.

46. Mukhopadhyay, D.; Dasso, M. Modification in reverse: the SUMO proteases. *Trends in Biochemical Sciences* **2007**, *32*, 286–295. https://doi.org/10.1016/j.tibs.2007.05.002.
47. Xiao, W.; Wang, R.S.; Handy, D.E.; Loscalzo, J. NAD(H) and NADP(H) Redox Couples and Cellular Energy Metabolism. *Antioxidants & Redox Signaling* **2018**, *28*, 251–272, [https://journals.sagepub.com/doi/pdf/10.1089/ars.2017.7216]. https://doi.org/10.1089/ars.2017.7216.
48. Böttinger, L.; Mårtensson, C.U.; Song, J.; Zufall, N.; Wiedemann, N.; Becker, T. Respiratory chain supercomplexes associate with the cysteine desulfurase complex of the iron–sulfur cluster assembly machinery. *Molecular Biology of the Cell* **2018**, *29*, 776–785, [https://doi.org/10.1091/mbc.E17-09-0555]. PMID: 29386296, https://doi.org/10.1091/mbc.E17-09-0555.
49. Stehling, O.; Lill, R. The Role of Mitochondria in Cellular Iron–Sulfur Protein Biogenesis: Mechanisms, Connected Processes, and Diseases. *Cold Spring Harbor Perspectives in Biology* **2013**, *5*, [http://cshperspectives.cshlp.org/content/5/8/a011312.full.pdf+html]. https://doi.org/10.1101/cshperspect.a011312.
50. Braymer, J.J.; Lill, R. Iron–sulfur cluster biogenesis and trafficking in mitochondria. *Journal of Biological Chemistry* **2017**, *292*, 12754–12763. https://doi.org/10.1074/jbc.R117.787101.
51. Shi, R.; Hou, W.; Wang, Z.Q.; Xu, X. Biogenesis of Iron–Sulfur Clusters and Their Role in DNA Metabolism. *Frontiers in Cell and Developmental Biology* **2021**, *Volume 9 - 2021*. https://doi.org/10.3389/fcell.2021.735678.
52. Petronek, M.S.; Allen, B.G. Maintenance of genome integrity by the late-acting cytoplasmic iron-sulfur assembly (CIA) complex. *Frontiers in Genetics* **2023**, *Volume 14 - 2023*. https://doi.org/10.3389/fgene.2023.1152398.
53. Gan, W.; Dai, X.; Dai, X.; Xie, J.; Yin, S.; Zhu, J.; Wang, C.; Liu, Y.; Guo, J.; Wang, M.; et al. LATS suppresses mTORC1 activity to directly coordinate Hippo and mTORC1 pathways in growth control. *Nature Cell Biology* **2020**, *22*, 246–256. https://doi.org/10.1038/s41556-020-0463-6.
54. Tsai, C.R.; Martin, J.F. Chapter Three - Hippo signaling in cardiac fibroblasts during development, tissue repair, and fibrosis. In *Cell Signaling Pathways in Development*; Soriano, P.M., Ed.; Academic Press, 2022; Vol. 149, *Current Topics in Developmental Biology*, pp. 91–121. https://doi.org/https://doi.org/10.1016/bs.ctdb.2022.02.010.
55. Gan, W.; Dai, X.; Dai, X.; Xie, J.; Yin, S.; Zhu, J.; Wang, C.; Liu, Y.; Guo, J.; Wang, M.; et al. LATS suppresses mTORC1 activity to directly coordinate Hippo and mTORC1 pathways in growth control. *Nature Cell Biology* **2020**, *22*, 246–256. https://doi.org/10.1038/s41556-020-0463-6.
56. Masri, S.; Cervantes, M.; Sassone-Corsi, P. The circadian clock and cell cycle: interconnected biological circuits. *Current Opinion in Cell Biology* **2013**, *25*, 730–734. Cell cycle, differentiation and disease, https://doi.org/https://doi.org/10.1016/j.ceb.2013.07.013.
57. Bevinakoppamath, S.; Ramachandra, S.C.; Yadav, A.K.; Basavaraj, V.; Vishwanath, P.; Prashant, A. Understanding the Emerging Link Between Circadian Rhythm, Nrf2 Pathway, and Breast Cancer to Overcome Drug Resistance. *Frontiers in Pharmacology* **2022**, *Volume 12 - 2021*. https://doi.org/10.3389/fphar.2021.719631.
58. Liu, J.; Jiang, Z.; Zha, J.; Lin, Q.; He, W. Crosstalk between the circadian clock, intestinal stem cell niche, and epithelial cell fate decision. *Genes & Diseases* **2025**, *12*, 101650. https://doi.org/https://doi.org/10.1016/j.gendis.2025.101650.
59. Bernasocchi, T.; Mostoslavsky, R. Subcellular one carbon metabolism in cancer, aging and epigenetics. *Frontiers in Epigenetics and Epigenomics* **2024**, *Volume 2 - 2024*. https://doi.org/10.3389/freae.2024.1451971.
60. Serefidou, M.; Venkatasubramani, A.V.; Imhof, A. The Impact of One Carbon Metabolism on Histone Methylation. *Frontiers in Genetics* **2019**, *Volume 10 - 2019*. https://doi.org/10.3389/fgene.2019.00764.
61. Stoccoro, A.; Lari, M.; Migliore, L.; Coppedè, F. Associations between Circulating Biomarkers of One-Carbon Metabolism and Mitochondrial D-Loop Region Methylation Levels. *Epigenomes* **2024**, *8*. https://doi.org/10.3390/epigenomes8040038.
62. Moretton, A.; Loizou, J.I. Interplay between Cellular Metabolism and the DNA Damage Response in Cancer. *Cancers* **2020**, *12*. https://doi.org/10.3390/cancers12082051.

63. Simpson-Lavy, K.J.; Bronstein, A.; Kupiec, M.; Johnston, M. Cross-Talk between Carbon Metabolism and the DNA Damage Response in S. cerevisiae. *Cell Reports* **2015**, *12*, 1865–1875. https://doi.org/10.1016/j.celrep.2015.08.025.
64. Borkúti, P.; Kristó, I.; Szabó, A.; Kovács, Z.; Vilmos, P. FERM domain–containing proteins are active components of the cell nucleus. *Life Science Alliance* **2024**, *7*, [https://www.life-science-alliance.org/content/7/4/e202302489.full.pdf]. https://doi.org/10.26508/lsa.202302489.
65. Ling, C.; Zheng, Y.; Yin, F.; Yu, J.; Huang, J.; Hong, Y.; Wu, S.; Pan, D. The apical transmembrane protein Crumbs functions as a tumor suppressor that regulates Hippo signaling by binding to Expanded. *Proceedings of the National Academy of Sciences* **2010**, *107*, 10532–10537, [https://www.pnas.org/doi/pdf/10.1073/pnas.1004279107]. https://doi.org/10.1073/pnas.1004279107.
66. Wang, S.; Zhou, L.; Ling, L.; Meng, X.; Chu, F.; Zhang, S.; Zhou, F. The Crosstalk Between Hippo-YAP Pathway and Innate Immunity. *Frontiers in Immunology* **2020**, *Volume 11 - 2020*. https://doi.org/10.3389/fimmu.2020.00323.
67. Dupont, S.; Morsut, L.; Aragona, M.; Enzo, E.; Giulitti, S.; Cordenonsi, M.; Zanconato, F.; Le Digabel, J.; Forcato, M.; Bicciato, S.; et al. Role of YAP/TAZ in mechanotransduction. *Nature* **2011**, *474*, 179–183. https://doi.org/10.1038/nature10137.
68. Mascanzoni, F.; Ayala, I.; Colanzi, A. Organelle Inheritance Control of Mitotic Entry and Progression: Implications for Tissue Homeostasis and Disease. *Frontiers in Cell and Developmental Biology* **2019**, *Volume 7 - 2019*. https://doi.org/10.3389/fcell.2019.00133.
69. Jesch, S.A.; Linstedt, A.D. The Golgi and Endoplasmic Reticulum Remain Independent during Mitosis in HeLa Cells. *Molecular Biology of the Cell* **1998**, *9*, 623–635, [https://doi.org/10.1091/mbc.9.3.623]. PMID: 9487131, https://doi.org/10.1091/mbc.9.3.623.
70. Iannitti, R.; Mascanzoni, F.; Colanzi, A.; Spano, D. The role of Golgi complex proteins in cell division and consequences of their dysregulation. *Frontiers in Cell and Developmental Biology* **2025**, *Volume 12 - 2024*. https://doi.org/10.3389/fcell.2024.1513472.
71. Woodward, A.M.; Göhler, T.; Luciani, M.G.; Oehlmann, M.; Ge, X.; Gartner, A.; Jackson, D.A.; Blow, J.J. Excess Mcm2–7 license dormant origins of replication that can be used under conditions of replicative stress. *Journal of Cell Biology* **2006**, *173*, 673–683, [https://rupress.org/jcb/article-pdf/173/5/673/1874564/673.pdf]. https://doi.org/10.1083/jcb.200602108.
72. Kolobynina, K.G.; Rapp, A.; Cardoso, M.C. Chromatin Ubiquitination Guides DNA Double Strand Break Signaling and Repair. *Frontiers in Cell and Developmental Biology* **2022**, *Volume 10 - 2022*. https://doi.org/10.3389/fcell.2022.928113.
73. Bassermann, F.; Eichner, R.; Pagano, M. The ubiquitin proteasome system — Implications for cell cycle control and the targeted treatment of cancer. *Biochimica et Biophysica Acta (BBA) - Molecular Cell Research* **2014**, *1843*, 150–162. Ubiquitin-Proteasome System, https://doi.org/https://doi.org/10.1016/j.bbamcr.2013.02.028.
74. Chang, Y.C.; Oram, M.K.; Bielinsky, A.K. SUMO-Targeted Ubiquitin Ligases and Their Functions in Maintaining Genome Stability. *International Journal of Molecular Sciences* **2021**, *22*. https://doi.org/10.3390/ijms22105391.
