## Supplementary material for "Novel 4D tensor decomposition-based approach integrating tri-omics profiling data can identify functionally relevant gene clusters": Supplementary_Table.pdf

Table S1: Factor Data Table of MOFA+

| | mRNA | trans | protein |
|---|---|---|---|
| Factor1 | $7.53 \times 10^{-2}$ | $4.97 \times 10^{1}$ | $2.10 \times 10^{-2}$ |
| Factor2 | $1.37 \times 10^{1}$ | $8.34 \times 10^{0}$ | $7.82 \times 10^{-1}$ |
| Factor3 | $8.31 \times 10^{0}$ | $5.75 \times 10^{-2}$ | $5.19 \times 10^{-2}$ |
| Factor4 | $4.01 \times 10^{0}$ | $3.16 \times 10^{0}$ | $1.78 \times 10^{-1}$ |
| Factor5 | $3.06 \times 10^{0}$ | $1.87 \times 10^{0}$ | $2.03 \times 10^{-1}$ |
| Factor6 | $1.70 \times 10^{0}$ | $9.21 \times 10^{-1}$ | $4.09 \times 10^{-3}$ |
| Factor7 | $1.38 \times 10^{0}$ | $3.18 \times 10^{-2}$ | $2.20 \times 10^{-3}$ |
